# Early Neutrophil Sensing Shapes the Innate Response to Preneoplastic Mammary Cells

**DOI:** 10.1101/2025.01.17.633444

**Authors:** Antoine Debiesse, Manuela Pereira Abrantes, Lyvia Moudombi, Aurélien Voissière, Leïla Lefèvre, Pacôme Lecot, Céline Rodriguez, Marine Malfroy, Iryna Moskalevska, Hélène Vanacker, Margaux Hubert, Albert Ndour, Roxane Pommier, Cassandra Assaf, Pierre Duplouye, Emmanuelle Jacquet, Marie Laurent, Sara Belluco, Nadège Goutagny, Anthony Ferrari, Nathalie Bendriss-Vermare, Virginie Petrilli, Julien Cherfils-Vicini, Christophe Caux, Marie-Cécile Michallet

**Affiliations:** Centre de Recherche en Cancérologie de Lyon (CRCL), Centre Léon Bérard, Université de Lyon, Université Claude Bernard Lyon 1, INSERM 1052, CNRS 5286, 69008 Lyon, France; Bayer HealthCare SAS, Medical Affairs, Lille 59000, France; Paris Centre University Hospital Infectious Diseases Stewardship team, AP-HP, Paris, France; Boiron France, Messimy, France; Laboratory for Immunotherapy of Cancer of Lyon (LICL), Centre de Recherche en Cancérologie de Lyon, Centre Léon Bérard, Lyon, France; Etablissement Français du Sang, Besançon, France; Université ColJte d’Azur, Centre National de la Recherche Scientifique (CNRS) UMR7284, Institut National de la Santé et de la Recherche Médicale (INSERM) U1081, Institute for Research on Cancer and Aging, Nice (IRCAN), Adipo-Cible Research Study Group, Nice 06107, France; Department of clinical oncology, Centre Léon Bérard, Lyon, France; Sanofi, Marcy L’Etoile, France; Synergie Lyon Cancer, ‘Gilles Thomas’ Bioinformatics Platform, Lyon, France; ADlab, Centre de recherche du CHU de Québec - Université Laval 2705 boul. Laurier, R-4773; Department of clinical oncology, CHU Grenoble Alpes, Grenoble, France; VetAgroSup, Marcy L’Etoile, France; Biomérieux, Marcy L’Etoile, France; Institut Hospitalo-Universitaire (IHU) RESPIRera and FHU OncoAge, CHU Nice, France

**Keywords:** Preneoplasia, Unfolded Protein Response, Immunosurveillance, Neutrophils, Breast Cancer

## Abstract

Breast cancer (BC) is the leading cause of cancer-related death in women. However, early detection of BC remains a major clinical challenge and represents a significant obstacle to effective prevention. To improve early clinical management, a deeper understanding of the preneoplastic immune microenvironment of BC is crucial. Among innate immune populations, neutrophils have emerged as important modulators of tumor development, but their role during the initiation of BC remains poorly understood. By integrating depletion experiments with transcriptomic profiling of sorted preneoplatic epithelial cells and neutrophils in spontaneous breast cancer mouse models, we observed that neutrophils contribute to tumor surveillance of preneoplastic stage with the activation of the unfolded protein response (UPR) in the preneoplastic epithelial compartment. To decipher the early anti-tumoral role of neutrophil, we developed an *in vitro* co-culture model of human mammary epithelial cells undergoing oncogenic stress with activation of the UPR (eHMEC), with human primary neutrophils. eHMEC display an immunoactive secretome as well as immunogenic membrane ligands, and neutrophils are the only immune cell population detecting eHMEC immunogenic signals leading to their recruitment, activation, production of reactive oxygen species and degranulation. Altogether, our work identifies for the first-time neutrophils as the earliest immune cell involved in immunosurveillance of preneoplastic BC epithelial cells, paving the way for potential therapeutic approaches targeting neutrophils to intercept early steps of BC tumorigenesis.

## INTRODUCTION

Breast cancer (BC) stands as the most prevalent cancer and a primary cause of cancer-related mortality among women globally^1^. BC develops over many years, progressing from preneoplastic lesions to invasive and metastatic disease^2–4^. Despite existing preventive measures including screening, chemoprevention, and emerging biological approaches, BC incidence continues to rise. Biological prevention, particularly strategies based on immunotherapy, remains limited by our incomplete understanding of how the immune system shapes the earliest steps of tumor development^5^.

The interaction between immune cells and preneoplastic or cancer cells is a permanent, dynamic, multifaceted, and evolving process. Numerous studies put their efforts to decipher the interplay between the immune system and clinically detectable established tumors. Within this immune compartment influencing BC progression^6^, the role of neutrophils, recognized as key players in combating pathogens, remains debated. Neutrophils, once thought to be a homogeneous population, are now recognized in both humans and mice as a heterogeneous compartment at the transcriptomic level^7–9^. Neutrophils exhibit a clear functional dichotomy, adopting either anti- or pro-tumoral roles recently associated to different stages of tumorigenesis. Indeed, neutrophils have been deeply characterized at late stages, where they exhibit pro-tumoral roles by promoting immunosuppression^10–13^ and cancer cell proliferation^14^, invasion and metastasis^15–17^, or angiogenesis^18^. In parallel, emerging studies have uncovered a previously underappreciated function for neutrophils at early cancer stages, where they contribute to early immune surveillance by promoting the detachment of transformed cells from the basal membrane^19^ and acting as antigen-presenting cells capable of stimulating T cell responses^20,21^. This functional shift from anti- to pro-tumoral state has been described as TGF-β dependent^22^.

In this study, we hypothesized that neutrophils play a key role in the surveillance of transformed mammary epithelial cells during the onset of BC. To investigate neutrophil activities at preneoplastic stage of BC tumorigenesis, we first used the MMTV-*Neu* breast cancer mouse model and developed an *in vitro* system of human mammary epithelial cells (HMEC). By neutrophil depletion in the MMTV-*Neu* model, we identified that neutrophils exert anti-tumor functions at early stages, before transitioning to pro-tumor activities at later stages. Transcriptomic analysis of sorted preneoplastic epithelial cells (pEpi) revealed an intrinsic activation of the unfolded protein response (UPR), validated in another public dataset of preneoplastic mammary cells, and an immunogenic signature characterized by the expression of *cxcl1*. Using an *in vitro* model of human mammary epithelial cells overexpressing HRas^G12V^ (eHMEC), we observed a similar activation of the UPR response, which regulates immunogenic features including a secretome and the expression of membrane ligands relevant to neutrophil. We next validated the specific neutrophil recruitment by eHMEC, followed by their activation, degranulation and reactive oxygen species (ROS) production. Altogether, our results show that neutrophils serve as the first line of defense during BC onset, due to their unique capacity to sense and be activated by stressed mammary epithelial cells.

## METHODS

### Mouse tumor models and experiments

#### MMTV-Neu mouse model

MMTV-NeuT transgenic mice were provided by Dr. Virginie Petrilli CRCL, and originally generated by Federica Cavallo University of Turin^23,24^ and bred under specific pathogen-free (SPF) conditions at the Small animal core facility/imaging platform (P-PAC platform, CRCL, Lyon, France). Neu+/− BALB/c males were crossed with wild-type BALB/c females (Charles River Laboratories). As described in the literature, the preneoplastic stage of tumor development, characterized by atypical mammary hyperplasia, occurs between the 6th and 10th weeks in mice expressing the Neu oncogene (NeuT+)^25^. Late stage, characterized by invasive carcinoma, occurs between the 20th and 25th weeks in mice expressing the Neu oncogene. Experiments were approved by the local and the French Ethical Committee for Animal Experimentation (#30253).

#### BLG-Cre, BRCA1^f/f^, p53^+/−^ mouse model

Cre recombinase expression is driven by the β-lactoglobulin (BLG) promoter, and exons 22-24 of BRCA1 are floxed, while one allele of Trp53 is mutated^26^. To initiate tumorigenesis through conditional BRCA1 deletion in the secretory epithelial cells of mammary glands, female BLG-Cre, BRCA1^f/f^, p53^+/−^mice were mated with male BRCA1^f/f^, p53^+/−^ mice. To ensure 100% penetrance, female mice underwent two cycles of gestation and lactation, developing spontaneous and sporadic mammary tumors 18 to 56 weeks after Cre activation, with a median onset at 36 weeks. For control mice, female BRCA1^f/f^, p53^+/+^ mice were mated with male BLG-Cre, BRCA1^f/f^, p53^+/−^ mice and also underwent two cycles of gestation and lactation to maintain comparable physiological conditions. At 36 weeks post-first mating, tumor-free mammary glands from female BLG-Cre, BRCA1^f/f^, p53^+/−^ mice (representing the early stage of tumor development) and healthy mammary glands from control mice were harvested, and cell suspensions were prepared for staining with a specific surface marker panel (**Table S2a**). Experiments were approved by the local and the French Ethical Committee for Animal Experimentation (#38957).

#### Anesthesia and Imaging Procedure

Mice were anesthetized by inhalation of 4% isoflurane (Forene®, Abbvie) for 5 minutes using a calibrated vaporizer. Anesthesia was subsequently maintained with 3% isoflurane throughout the imaging procedure. Physiological parameters, including respiratory rate, heart rate, and body temperature, were continuously monitored to ensure animal welfare.

The imaging area, extending from the cervical to the abdomen, was prepared by hair removal using standard shaving and depilation procedures.

Mice were then positioned on the Vevo LAZR Multimodal Imaging System (FUJIFILM VisualSonics Inc.) for ultrasound acquisition using a coupling medium (Aquasonic Clear® 100 Transmission Gel, Ref. 03-50). High-resolution images and three-dimensional scans were acquired for both left and right upper mammary glands. Tumor volume was quantified by 3D reconstruction using Vevo LAB software (FUJIFILM VisualSonics Inc., v3.0.0).

#### Histopathological staining

Organs were formalin-fixed, paraffin-embedded, and processed for hematoxylin and eosin staining (Research pathology platform East (PAR), CRCL, Lyon, France). The pathological examination of the slides was performed by Dr. Sara Belluco, a veterinary pathologist from VetAgro Sup (Ecole Vétérinaire Nationale, Lyon, France). All glands analyzed at the preneoplastic stage were first evaluated histopathologically to confirm the complete absence of invasive carcinoma; samples showing any invasion were excluded from analysis.

#### Blood Cell Isolation

Peripheral blood was collected three days prior to organ harvesting to permit restoration of granulopoiesis homeostasis. A drop of tetracaine (single-dose ophthalmic solution) was applied to the mouse’s eye prior to the collection of approximately 200 µL of blood via retro-orbital sampling and immediately mixed with 15 µL of 0.5 M EDTA (Sigma-Aldrich). A volume of 100 µL of whole blood was diluted in 1 mL of 1X PBS (Gibco, #14200-067) containing 0.004 M EDTA, followed by centrifugation (10 min, 1200 rpm, room temperature). The cell pellet was resuspended in 1 mL of 1X PBS, incubated with 10 mL of 1X Pharm Lyse buffer (BD, #555899) for 2–5 minutes at room temperature in the dark, with gentle mixing performed twice during incubation. Following a wash step with 40 mL of 1X PBS, samples were centrifuged. The final cell pellet was resuspended and transferred to a 96-well plate for subsequent staining.

#### Generation of cell suspensions from mammary and lymph nodes tissues

Mice were euthanized by cervical dislocation and mammary and lymph nodes tissues were collected, weighed, minced, and digested with 1mg/mL Collagenase IA and 0,02mg/mL DNase (Sigma, #C2674 and #D4513) for 30’ at 37°C. The digested tissues were filtered on 70µm and 30µm cell strainers, centrifuged (300g, 10’, 10°C). Cells were resuspended in RPMI 1640 (Gibco, #72400-021) supplemented with 100U/mL of penicillin and 100µg/mL of streptomycin (Gibco, #15140-122) and 10% FBS (Eurobio, #CVFSVF00-01) for further experiments.

#### Generation of cell suspensions from bone marrow and spleen

Mice were euthanized by cervical dislocation to collect femoral bone marrow, spleen and lymph node. Bone marrow was flushed with 4–5 mL of complete RPMI 1640 medium (supplemented with GlutaMAX™, 2% FBS, 2 mM EDTA, and 1% penicillin/streptomycin), centrifuged (5 min, 1500 rpm, RT), and filtered sequentially through 70 µm and 30 µm strainers. Cells were counted with Türk’s blue and prepared for subsequent labeling and sorting.

Spleen was mechanically dissociated on a 70 µm cell strainer using a syringe plunger. Filter was rinsed three times with 10 mL of complete RPMI medium (RPMI – 10% FBS – 1% L-Glutamin – 1% Penicillin/Streptomycin). Cell suspension was centrifuged at 1500 rpm for 5 minutes at RT and resuspended in 5 mL of 1X PharmLyse solution (#555899, BD), followed by incubation at room temperature for 3–5 minutes. Subsequently, 20 mL of 1X PBS was added, and the suspension was centrifuged under the same conditions for an additional wash with 30 mL of 1X PBS. Finally, the pellet was resuspended in 2 to 10 mL of RPMIc for cell counting using Turk’s blue. After a final centrifugation, cell concentration was adjusted to 5 × 10^7 cells/mL using RPMIc. If necessary, pellets with lower concentrations were resuspended in 200 µL of RPMIc.

#### Short-term neutrophil depletion at late stage of tumorigenesis

Late-stage neutrophil depletion was performed by intraperitoneal injection of a depleting anti-Ly6G monoclonal antibody (clone 1A8, InVivoPlus anti-mouse Ly6G, #BP0075-1, BioXcell) in NeuT^+^ mice (n=14). Control NeuT^+^ mice (n=11) received an isotype control antibody (InVivoPlus rat IgG2a isotype control, #BP0089, BioXcell). Injections were administered three times per week (Monday, Wednesday, Friday) at a dose of 200_µg for three consecutive weeks, starting from the detection of tumors reaching 15–30□mm^3^ in volume (approximately at 15 weeks of age). To assess depletion efficiency, ∼100□µL of blood was collected retro-orbitally three hours prior the first injection.

#### Long-term neutrophil depletion at preneoplastic stage of tumorigenesis

Early-stage neutrophil depletion^27^ was induced using a combination of antibodies to enhance depletion over a 7-week period. NeuT⁺ mice received daily intraperitoneal injections (Monday to Saturday) of anti-Ly6G antibody (InVivoPlus anti-mouse Ly6G, #BP0075-1, BioXcell), while control NeuT+ mice were treated with the corresponding isotype control (InVivoPlus rat IgG2a isotype control, #BP0089, BioXcell). In addition, a secondary anti-rat kappa light chain antibody (InVivoMab anti-rat Kappa Light Chain, #BE0122, BioXcell) was administered every other day (Monday, Wednesday, Friday), two hours after the primary antibody. This protocol was maintained for 7 weeks starting at 4 to 6 weeks of age. Depletion efficiency was monitored weekly via submandibular blood collection (∼20□µL) performed prior to Monday injections.

For long-term neutrophil depletion, two cohorts were conducted with 5 and 4 mice in the control NeuT+ group, and 8 and 10 mice in the NeuT+ group, respectively.

#### Cell Sorting of Epithelial Cells from mammary glands of NeuT− or NeuT+ mice

Cells previously isolated from the mammary glands of NeuT− or NeuT+ mice at preneoplastic and late stages were incubated with FcBlock (Purified, Biolegend, #101301) for 15’ at 4°C in the dark, followed by staining with a mix of fluorescent-conjugated antibodies (**Table S2b**) to identify epithelial cells (Lin^−^CD45^−^ EpCAM^+^), incubating for 30’ at 4°C in the dark. Cells were washed twice with FACS buffer (PBS1x + 2mM EDTA, 3% FBS). After the final centrifugation (5‘, 1500rpm, 4°C), cells were resuspended and filtered in a 1X DAPI solution diluted in Pre-Sort FACS buffer (BD, #563503) at a concentration of 15.10^6^ cells/ml. Epithelial cells (5000 cells) were sorted by flow cytometry (BD FACS Aria™ II) into 300µl of TCL lysis buffer (Qiagen, #1031576) supplemented with 1% β-mercaptoethanol (Sigma, #M6250). Tubes were vortexed briefly, immediately snap-frozen in liquid nitrogen, and stored at −80°C.

#### Cell Sorting of Neutrophils from mammary glands of NeuT− or NeuT+ mice at preneoplastic and late stages

Cells previously isolated from the mammary glands of NeuT− or NeuT+ mice at preneoplastic and late stages were enriched for CD45⁺ cells prior to immunofluorescent staining and FACS-sorting using EasySep mouse CD45 positive selection kit (#18945, StemCell Technologies) according to the manufacturer’s instructions.

Then, cells were incubated with FcBlock (Purified, Biolegend, ref#101301) for 15’ at 4°C in the dark, followed by staining with a mix of fluorescent-conjugated antibodies (**Table S2c**) to identify neutrophils (Lin− CD45.2^+^ Ly6G^hi^ Ly6C^−/int^), incubating for 30’ at 4°C in the dark. Cells were washed twice with FACS buffer (PBS1x + 2mM EDTA, 3% FBS). After the final centrifugation (5‘, 1500rpm, 4°C), cells were resuspended and filtered in a 1X DAPI solution diluted in Pre-Sort FACS buffer (BD, #563503) at a concentration of 15.10^6^ cells/ml. Neutrophils (1000 cells) were sorted by flow cytometry (BD FACS Aria™ II) into 300µl of TCL lysis buffer (Qiagen, #1031576) supplemented with 1% β-mercaptoethanol (Sigma, #M6250). Tubes were vortexed briefly, immediately snap-frozen in liquid nitrogen, and stored at −80°C.

#### Library Preparation and RNA-Sequencing

Total RNA was extracted from 1000 neutrophils and 5000 epithelial cells using the Single Cell RNA Purification Kit (Norgen, ref#51800), according to manufacturer’s instructions, with an additional RNAse-free DNAse treatment (Qiagen, #79254). RNA quality (RNQ and DV200) was assessed using the Ultra Sentivity RNA Kit (Agilent, #FP-1201-0275) and the FemtoPulse system (Agilent) at the Gentyane Platform (Clermont-Ferrand, France). A pre-treatment with DNase using the Heat&Run gDNA Removal Kit (ArcticZymes Technologies) was performed prior to reverse transcription to prevent genomic DNA contamination. Following to manufacturer’s instructions, libraries were prepared with the SMART-Seq Stranded Mammalian Single Cell Kit (Takara/Clontech, #634442) and sequenced on a NovaSeq Illumina platform (Cancer Genomics Platform, CRCL, Lyon, France) with a paired-end mode and a targeted depth of 32M reads/sample.

#### RNA-seq data processing

RNA libraries were prepared with the TrueSeq poly-A+ kit and sequenced on an Illumina NovaSeq 6000 sequencing machin e. For careful quality controls, raw data were aligned on the human genome (GRCh38) with STAR (v2.7.8a)^28^, with the annotation of known genes from gencode v37. RNA quality controls metrics were computed using RSeQC (v4.0.0)^29^. Gene expression was quantified with Salmon (v1.5)^30^ on the raw sequencing reads, using the annotation from gencode v38 as index. Further gene expression analyses were restricted to protein-coding genes using annotables R package (v.0.2.0) (protein coding, immunoglobulins (IG) and T cell receptors (TR) biotypes) (https://github.com/stephenturner/annotables). Starting from log2-transformed, TPM-normalized data, unsupervised analyses (Principal Component Analysis, PCA, and hierarchical clustering) were performed on the top 10% most variable protein-coding genes from the cohorts of interest (infiltrating neutrophils or epithelial cells). PCAs were performed with the stats R package (v4.4.1) and visualized using factoextra (v1.0.7) (https://github.com/kassambara/factoextra). Hierarchical clustering and heatmaps generation were carried out with ComplexHeatmap (v2.20.0) using the Ward.D linkage method and Euclidean distance^31^. Pathway lists originated from MSigDB (Molecular Signatures Database) and were obtained through msgidb (v1.12) R package (v.1.12.0)^32,33^. Single sample GSEA (ssGSEA) scores were computed on TPM normalized data through gsva R package (v1.52.3)^34^.

Differential expression analyses were performed using the R package DESeq2 (v1.44.0)^35^, with Wald test and apeglm shrinkage estimator (v1.24.0)^36^. Volcano plots were generated with the ggplot2 R package (v3.5.1)^37^. (**Table S3a,b**)

#### Characterization of the immune landscape by flow cytometry

After generation of cell suspensions from mammary tissues as described above, immunophenotyping of the immune compartment was performed using a panel of 29 markers (**Table S2d**) designed by the Flow Cytometry Core Facility of the CRCL. In collaboration with the Core Facility, we carried out stainings, data acquisition and processing, gating, and downstream analyses.

#### Characterization of the neutrophils infiltrate by Multiplex Immunofluorescence (mIF)

Fully automated three-colors multiplex immuno-fluorescence was performed with the OPAL system (Akoya Biosciences) using the BOND RX stainer (Leica Biosystems). Four-micrometer sections from formaldehyde-fixed and paraffin-embedded (FFPE) murine mammary glands were deparaffinized, rehydrated and antigen retrieval treatment was performed using ER1 (for CD45 and EpCAM) or ER2 (for Ly6G) buffer for 20’ at 95°C. The sections were sequentially stained with each primary antibody, followed by OPAL-HRP secondary antibody incubation then revealed with tyramide signal amplification and OPAL fluorophore, in the following order: EpCAM (polyclonal; #ab71916, Abcam; 1/500 dilution)/OPAL 690; Ly6G (clone E6Z1T; #87048S, CST; 1/300 dilution)/OPAL 570; CD45 (clone D3F8Q; #70257, CST; 1/300 dilution)/OPAL 520. The sections were then counterstained with spectral DAPI (Akoya Biosciences) and mounted with a coverslip (**Table S4**). Whole slides were imaged at a 20x magnification using the Vectra Polaris multispectral scanner (Akoya Biosciences) and digital images were visualized with the Phenochart viewer (Akoya Biosciences) and unmixed using the spectral library from the software. Quantification of Ly6G labeling was performed using Halo software (Indica Labs).

#### ElectroChemiLuminescence Immuno Assay (ECLIA)

Cytokine quantification in supernatants of dilaceration of mammary glands. The following cytokines were quantified in supernatants of dilaceration of mammary glands from NeuT− or NeuT+ mice at early and late stage using MSD technology according to the U-plex protocol from Mesoscale Discovery (MSD) and separated into two kits: TGF-β1, TGF-β2, TGF-β3 and KC/GRO (**Table S5a**). Supernatants were tested without dilution before quantification.

#### Analysis of Publicly scRNA-seq dataset from BRCA1-mutated mammary tumor model

Publicly available scRNA-seq dataset from the BRCA1-mutated mouse mammary tumor model (BLG-CRE, BRCA^f/f^, p53^+/−^)^38^ was utilized to characterized ER stress and UPR response on epithelial cells during tumor progression. In this study, the authors sorted live cells from mammary glands at different stages during tumor development and performed scRNA-seq using 10X Genomics technology. Dataset processing and analyses were performed using Seurat (version 5.0). After normalization (LogNormalize) and scaling, principal component analysis (PCA) was performed on non-tumor epithelial cells following a pseudo-bulk approach, where gene expression of individual cells within each sample was aggregated. Based on the PCA results, samples were stratified into WT-like and 2 pre-cancer stages : Stage-I and Stage-II. Gene signature scores of ER stress and UPR pathways from REACTOME, HALLMARKS and Gene Ontology (GO) databases (**Table S3a**), were calculated for each cell using Ucell package (version 2.10.1) and the *AddModuleScore_UCell* function of Seurat.

### Human models and experiments

#### Human Mammary Epithelial Cell Line Generation

Human mammary epithelial cells (HMEC - Promocell, #C-12650) were immortalized with by transduction with a retroviral vector pBABE-puro containing the Human Telomerase Reverse Transcriptase (hTERT - Addgene). HMEC-hTERT cells then transduced with a lentiviral plasmid/vector pLVUT’ for doxycycline-inducible HRas^G12V^ (HMEC-HRas) of Green Fluorescence Protein (HMEC-GFP) expression (tet ON system - Trono/Aebischer’s lab. Cells were cultured in Mammary Epithelium Cell Growth Medium (MECGm – Promocell, #CC-5012) supplemented with 100U/mL of penicillin and 100µg/mL of streptomycin (Gibco, #15140-122). Upon reaching 75% confluence, cells were washed with PBS1x (Gibco, ref#11540486), trypsinized with Trypsin/EDTA (0.25 mg/mL - Lonza, #CC-5012) for 20’ at 37°C, and neutralized with Trypsin Neutralizing Solution (TNS - Lonza, #CC-5002). Cell suspension was centrifuged (5’, 1400rpm, room temperature (RT)), and the pellet was re-suspended in 37°C MECGm before plating.

#### In vitro HRas^G12V^ and GFP Induction Assay

At day −1 (d-1), cells were plated at 1% of confluency in culture dishes or culture plates. At d0, culture medium was replaced by MECGm supplemented or not with doxycycline (1µg/mL – Sigma, #D9891-1G) in order to induce or not HRas^G12V^ or GFP. Supplemented mediums were then daily refreshed.

#### Human Immune cells Isolation

Immune cells were isolated from healthy donor blood samples of the “Etablissement Français du Sang” (Lyon) according manufacturer’s instruction. Briefly, to obtain total immune cells, blood was lysed with 1x lysing buffer (4,5mL of blood for 45mL of lysing buffer, BD Pharm Lyse – BD, #555899). To isolate peripheral blood mononuclear cells (PBMCs), the standard Ficoll method (Eurobio, #CMSMSL01-01) was used. For neutrophil isolation, the EasySep™ Direct Human Neutrophil Isolation Kit (Stemcell, #19666) was employed.

#### Morphology Assessment

Cells were imaged daily using an Axiovert25 microscope (Zeiss) with 20x magnification and analyzed using Zen software.

#### SA-**β**-galactosidase Activity

β-galactosidase staining was performed using the Cell Signaling β-galactosidase staining kit (Cell signaling, #9860). Cells were fixed and stained with X-Gal substrate (1mg/mL, pH6) overnight at 37°C in a dry incubator.

#### Microarray Analyses

Microarrays analyses were performed to identify differentially expressed genes/pathways between d3 (eHMEC of kinetics - n=4) and d5 (sHMEC of kinetics - n=3) and their control. Briefly, at d3 or d5 of the kinetics, supernatants were removed, and cells were washed with PBS1x. Then, cells were lysed directly in plate, in 350μl of RLT buffer (Qiagen, #79216). Lysates were vortexed, snap frozen and conserved at – 80°C. RNA extraction, reverse transcription and microarray hybridization on Affymetrix GeneChip Human PrimeView arrays (Affymetrix). Microarray data analysis was conducted in R (version 4.3.3). Raw probe intensities were quantile normalized and log2-transformed using the Robust Multi-array Averaging (RMA) method available in the affy package (v. 1.80.0). We identified Differentially expressed genes (DEG) between eHMEC and their control by using DESeq2 (v. 1.42.1) with an adjusted p-value (adj.P.Val) < 0.05 and a log2 fold change > 0.58. Their locations were determined by using the database Uniprot (https://www.uniprot.org/). Single sample Gene set enrichment analyses (ssGSEA) were performed by using the GSVA package (v. 1.50.5) and the Molecular Signatures Database into Human collection (Human MSigDB v2024.1.Hs). Used genesets are presented in **Table S3c**. The statistical significance of ssGSEA score was addressed with two-tailed Wilcoxon test, with ⍰=0,05.

#### Immunoblotting

HMEC-HRas were lysed on ice in Laemmli 2x buffer (0.125M Tris HCl pH 6.8 (Euromedex, #EU0011; Fisher, #H/1150/PB15), 2% SDS (Euromedex, #EU0460), 0.5M DTT (Euromedex, #EU0006-D)). Protein quantification was performed using Bradford reagent (Bio-Rad, #5000006) with Bovine Serum Albumin standards (Sigma, #A9647). For each condition, 20μg of denatured protein was resolved on 8 or 12% SDS-PAGE and transferred to nitrocellulose membranes using the Trans-Blot® Turbo Transfer System (Bio-Rad, #17001918). Membranes were blocked in TBS-Tween 5% BSA, probed with primary antibodies (**Table S6**) overnight at 4°C, washed, and incubated with HRPconjugated secondary antibodies. Signals were detected using Clarity™ Western ECL substrate (BioRad, #170-5061) and the ChemiDoc Imaging System (Bio-Rad). Results were analyzed with Image Lab software (Bio-Rad).

#### RNA Extraction and RT-qPCR

Total RNA was extracted using the Nucleospin RNA kit (Macherey Nagel, #740955) according to the manufacturer’s instructions. RNA was quantified by NanoDropTM One (Thermo Scientific), and retrotranscription of 1000ng of RNA was performed using the iScript kit (Bio-Rad, #1725038). qPCR was carried out using a CFX Connect™ Real-Time System (Bio-Rad) with SYBR Green (Bio-Rad, #1725271) and 10µM primers (**Table S7**) for a total of 40 cycles, as follows: 95°C for 15sec, 60°C for 20sec, and 95°C for 15sec. Results were represented as mean ratio of the gene of interest on the housekeeping gene (GUS), and normalize as fold of the corresponding “control” sample.

#### Enzyme Linked ImmunoSorbent Assay (ELISA)

ELISA kits (**Table S5b**) were used according to the manufacturer’s instructions. Briefly, ELISA plates (Nunc Maxisorp - Thermo Scientific, #442404) were coated with capture antibodies, blocked with PBS1X - 1% BSA, and incubated with cell line supernatants, standards, or controls in duplicates. After washing, HRP-conjugated detection antibodies were added, and 3,3’,5,5’-tetramethylbenzidine (TMB) enzymatic substrate (Interchim, #UP664781) was used for signal development. Optical density was read at 450nm with correction at 540–570nm using a Multiskan FC reader (Thermo Scientific).

#### Transwell Migration Assay

After isolation, immune cells, PBMCs and isolated neutrophils pellets were resuspended in MECGm at 5×10□cells/mL. Conditioned medium was collected from control HMEC and eHMEC wells on day 3 and normalized to the number of cells per well. A volume of 600□μL of the normalized medium was added to the lower compartment of Transwell plates. Then, 100□μL of immune cells, PBMCs or isolated neutrophils was added to the upper inserts with 3□μm pore size membranes (Corning, #003415). For assays involving total immune cells or PBMCs, cells were added to the upper inserts with 5□μm pore size membranes (Corning, #003421). All conditions were performed in duplicate and cells migrated for 3h at 37°C, 5% CO_2_. Immune cells were stained with a specific surface marker panel (**Table S2e-f**) and 20μL of Flow Count beads (Beckman Coulter, #7547053) were added before analysis.

#### p-STAT5 Activation

Total immune cells were resuspended in 1mL of conditioned medium (normalized to cell culture number) to achieve 2×10□ immune cells/mL. Neutrophils were incubated for 30’ at 37°C, 5% CO_2_, centrifuged (5’, 1500rpm, 37°C) and stained with specific surface marker panel (**Table S2g**).

#### ROS Production

Isolated neutrophils were resuspended in 1mL of conditioned supernatants (normalized to cell culture number) at 0,5×10□ neutrophils/mL and incubated for 12h at 37°C, 5% CO₂. Neutrophils were transferred to a 96-well V-bottom plate on ice, washed with PBS1x, and stained with Zombie NIR for viability. After two washes with FACS buffer (PBS1x + 2 mM EDTA, 3% FBS), pellets were resuspended in 100μL of ROS-detection solution (CM-H2DCFDA - Invitrogen, #C6827) and incubated for 20’ at RT, protected from light. Cells were washed and stained with a specific surface marker panel (**Table S2h**).

#### Co-culture Assay

At d2,5 cells were co-cultured with isolated neutrophils at a ratio of 1 HMEC to 5 neutrophils for 12h. After co-culture, cells were harvested, centrifuged (5’, 1500rpm, RT) and stained with a specific surface marker panel (**Table S2i**) to assess neutrophil activation and degranulation.

#### *In Vitro* Matrigel Assay

At d0, HMEC were resuspended in cold MECGm at 0,5×10_ cells/mL. A 100µL cell suspension was mixed with 500µl Matrigel growth factor reduced (Corning, #354230), with or without dox (1,7µg/mL), on ice. The mixture was gently agitated, incubated on ice for 1h, then drawn into a cold syringe and layered in a 12w plate. Plates were incubated at 37°C with 5% CO_2_ for 45’, followed by the addition of 1mL MECGm with or without dox (1µg/mL). Media were refreshed daily. At d3 or d5, supernatants were collected and matrigel with cells were recovered with 2mL cold MECGm, transferred to a 50mL tube, diluted with 28mL cold MECGm, and centrifuged twice (5‘, 1500 rpm, 4°C). Cells were treated with 10mL Trypsin-EDTA 0.05% (Gibco, #25300-054) for 20’ at 37°C with 5% CO_2_, followed by neutralization with 10mL MECGm. After centrifugation (5’, 1500rpm, RT), cells were counted and snap-frozen.

#### *In Vivo* Matrigel Plug Assay

Experiments were performed as previously described^39^, on female NMRI Nude mice (Charles River). A total of 50 000 HMEC were suspended in 100μl PBS1x and mixed with 400μL Matrigel growth factor reduced (Corning, #354230) with or without dox (1,7µg/mL), on ice. The mixture was immediatly injected subcutaneously into the back of mice under isoflurane anesthesia. After three days, mice were sacrificed, and Matrigel plugs were collected. Infiltrating cells were isolated by enzymatic dissociation using Dispase (Corning, #354235), Collagenase A (Roche, #10103586001), and DNAse I (Roche, #05952077103) for 30’ at 37°C and were stained with a specific surface marker panel (**Table S2j**).

#### Cytotoxicity assay

At d2,5, dox-treated or not cells were co-cultured with isolated neutrophils from healthy donor blood at a ratio of 1 eHMEC to 5, 10 or 20 neutrophils for 24h in flat-bottom 24-well plates. Neutrophils were subsequently removed by two washes with 250 µL of 1X PBS per well. Then, 500 µL per well of PrestoBlue reagent (diluted 1:10 in complete MECGm medium) was added in the plate, followed by incubation for 1 hour at 37°C in in a cell culture incubator, protected from direct light. Cell viability can be detected using a fluorescence-based microplate reader M1000 (Tecan) with fluorescence excitation wavelength of 545 nm and an emission of 585 nm. Background fluorescence was corrected by including control wells containing only cell culture medium (no cells) on each plate. Cell viability percentages (% viable cells) were normalized to control eHMEC or eHMEC without neutrophils.

#### Conventional Flow Cytometry

##### Extracellular Staining

Cell suspensions were plated in a 96-well V-bottom plate, centrifuged (5’ at 1400rpm, RT), and resuspended in 100μL to assess viability. After a 25’ incubation at 4°C in the dark, cells were washed twice with FACS buffer (PBS1x + 2 mM EDTA, 3% FBS) and incubated for 25’ at 4°C with a mix of a specific surface marker panel antibodies (Table S2). Following two washes with FACS buffer, cells were fixed with 4% formaldehyde (Sigma, #F8775-500ML) or Fix/Perm buffer (#00-5523-00, ebiosciences, for intracellular staining) for 20’ at 4°C. Cells were resuspended in 200μL of FACS buffer, transferred to Micronic tubes. Samples were analyzed on LSRFortessaTM (BD Biosciences) or Cytek® Aurora (Cytek Biosciences), and data were processed with FlowJo software.

##### Intracellular Staining

After extracellular staining, cells were washed with PBS1x and fixed in preheated BD Lyse/Fix Buffer (BD, ref# 558049) for 10 min at 37°C under 5% CO_2_. After washing with PBS1X, cells were permeabilized with pre-chilled Perm Buffer III (BD, ref# 558050) for 30’ on ice in the dark. Following washes, intracellular antibodies (Table S2) diluted in FACS buffer were added, and cells were incubated for 45’ in the dark. After a final wash, cells were resuspended in 200μL of FACS buffer, transferred to Micronic tubes, and analyzed using LSRFortessaTM (BD Biosciences) and FlowJo software.

#### Statistical Analysis

All data analyses were performed using Prism Version 10.3.1 (GraphPad Software). The number of independent experiments is indicated in the legends of each figure. First, a Shapiro-Wilk normality test was performed on all data sets. Then according to these results parametric (normality) and nonparametric (does not follow the normal law) tests were used. The statistical significance of RNA-seq data of MMTV-Neu mouse model was addressed with Welch’s t test with ⍰=0,05 (*p<0.05; **p<0.01; ***p<0.001; ****p<0.0001, ns p≥0.05). Late-stage depletion, public neutrophil signature enrichment, and KC/GRO quantification in the MMTV-*Neu* model were analyzed using the Mann–Whitney test (α = 0.05; *p < 0.05; **p < 0.01; ***p < 0.001; ****p < 0.0001; ns p ≥ 0.05). Immune and neutrophil infiltrates, gene expression analyses (*Tgfbr1*, *Siglecf*, *Cxcl1*), and TGF-β quantification in the MMTV-*Neu* model were analyzed using the Kruskal–Wallis test (α = 0.05; *p < 0.05; **p < 0.01; ***p < 0.001; **p < 0.0001; ns p ≥ 0.05). Immune cell enrichment based on FACS data in the MMTV-*Neu* model was analyzed using the Wilcoxon test (α = 0.05).The statistical significance of proliferation data was addressed with a Two-Way ANOVA test using Geisser-Greenhouse correction method, with IZ=0,05 (*p<0.05; **p<0.01, ***p<0.001; ****p <0.0001; non-significant results defined as p≥0.05 had ns). The statistical significance of RT-qPCR, ELISA, immunogenic membrane ligands staining, neutrophil migration and activation, matrigel assay and neutrophil infiltration in BLG-Cre, BRCA1^f/f^, p53^+/−^ mouse model data was addressed with a two-tailed Wilcoxon test (*p<0.05; **p<0.01, ***p<0.001; ****p <0.0001; non-significant results defined as p≥0.05 had ns). The statistical significance of microarray data was addressed with Mann-Whitney test or One-way ANOVA with IZ=0,05 (*p<0.05; **p<0.01; ***p<0.001; ****p<0.0001, ns p≥0.05). The statistical significance of data with HMEC treated or not with UPR inhibitors (ELISA, immunogenic membrane ligands) was addressed with RM-one-Way ANOVA test using Geisser-Greenhouse correction method, with IZ=0,05 (*p<0.05; **p<0.01; ***p<0.001; ****p<0.0001, ns p≥0.05).

## RESULTS

### Neutrophils display an anti-tumoral role at preneoplastic stage of BC *in vivo*

To study the role of neutrophils at preneoplastic stage, we used the spontaneous BC MMTV-*Neu* in which the MMTV promoter drives mammary-specific expression of HER2/*Neu* (herein referred to as NeuT+), predominantly within alveolar luminal epithelial cells, along with control mice lacking the oncogene expression (NeuT−) (**Fig S1a**). Using ultrasound imaging, we showed that NeuT+ mice develop tumors in an asynchronous manner, with some tumors initiating as early as 13 weeks and others appearing as late as 20 weeks of age (**Fig S1b-c**). To determine the preneoplastic stage, we performed histological analyses of mammary glands (MG). As expected, NeuT⁻ mice displayed normal mammary gland (hMG) development at both early and late timepoints. In contrast, NeuT⁺ mice showed a clear progression of lesions: at the early timepoint, 6–10 weeks, H&E staining revealed hyperplasia with dysplasia covering approximately 50% of the MG (preneoplastic stage, pMG). These preneoplastic lesions evolved into rare *in situ* carcinomas (<3%) and progressively into invasive carcinomas, which accounted for approximately 16% of the mammary gland by 10–12 weeks and steadily increased with age, reaching up to 95% by 20–25 weeks (tumor stage, tMG) (**Fig S1d–f**), as previously described^25^. Based on the asynchronous initiation and the heterogeneity observed, we defined the preneoplastic stage as occurring before 10 weeks of age in the absence of invasive carcinoma (**Fig S1d, S1f**). Regarding the immune infiltrate, pMG was comparable to that of early hMG, both in terms of immune cell proportions among living cells (**Fig S1g**) and overall composition (**Fig S1h**), in contrast with the marked increase and remodeling observed in tMG (**Fig S1g, S1i**). Focusing on neutrophils, we already detected their presence at the preneoplastic stage (**Fig S1h–i**). Although no statistically significant differences were observed in pMG compared to hMG, immunofluorescence staining of FFPE sections using the Ly6G marker revealed a trend toward increased PAN (Preneoplastic-Associated Neutrophils) within hyperplasia with dysplasia areas compared to healthy tissue (**Fig S1j**). Together, these data confirm that the MMTV-*Neu* model is relevant for investigating the impact of neutrophils during the preneoplastic stage.

To assess the role of PAN in preneoplastic stage, we performed neutrophil-specific depletion. At early timepoint, PAN depletion was also validated with a significant decrease of circulating neutrophils after 3 weeks, which persisted throughout the treatment (**Fig S2a**). Interestingly, PAN depletion was also effective in the pMG and lymph nodes, whereas the bone marrow and the spleen were not affected (**Fig S2b**). As expected, neutrophil-depleted and control isotype-treated NeuT+ mice developed tumors (**Fig S2c**) whereas PAN depletion accelerated tumor growth in almost half of the treated animals (8/18 mice) (**Fig 1a**). As expected, TAN (Tumor-Associated Neutrophils) exert a pro-tumoral role at the late stage of BC (**Fig S2d-e**). Taken together, our findings indicate that PAN contribute to the surveillance of preneoplastic cells.

**Figure 1:**
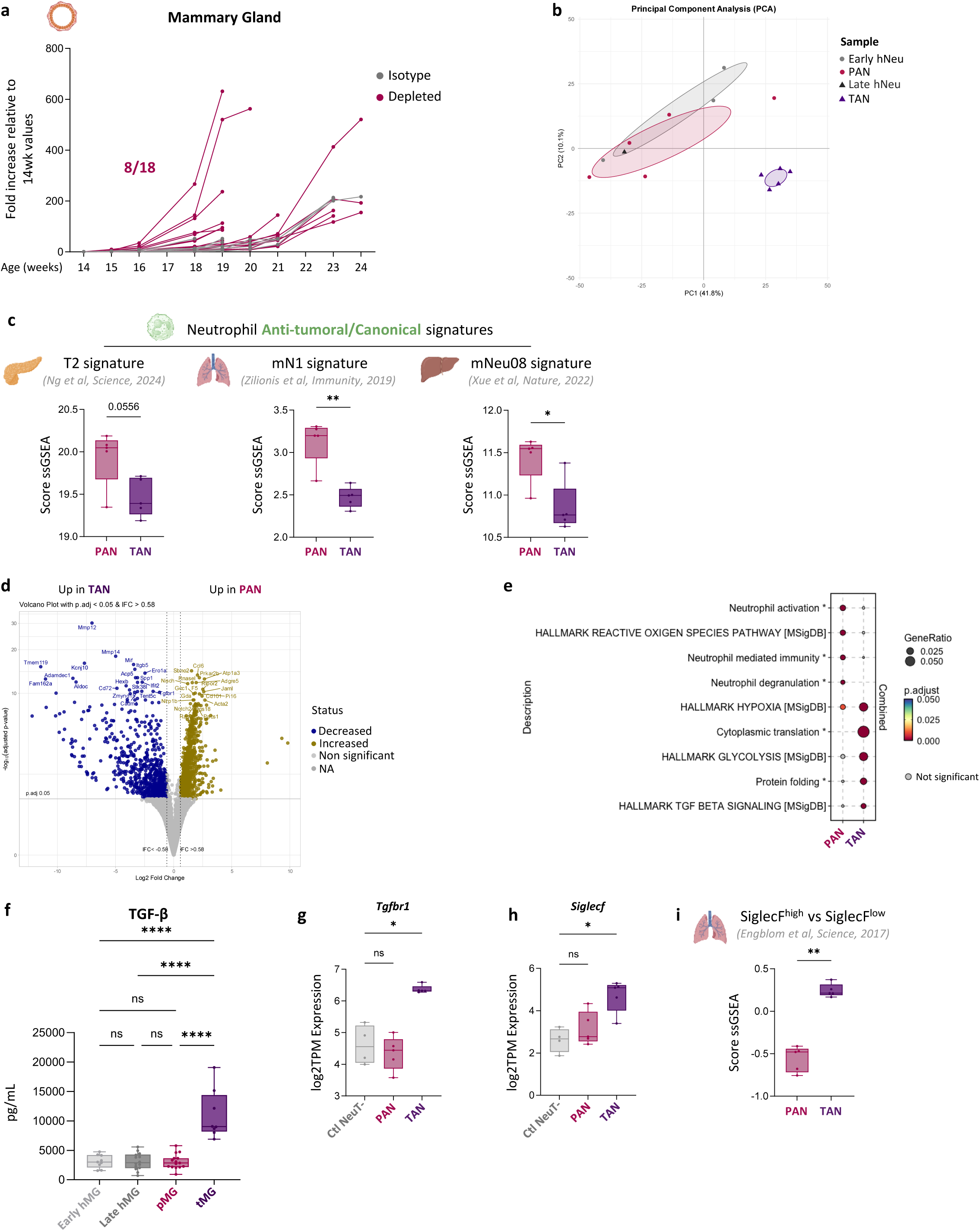
Preneoplastic-Associated Neutrophils are anti-tumoral but are reprogrammed at late stages. **a.** Monitoring of tumor growth across weeks of development after neutrophil depletion at the preneoplastic stage, expressed as fold increase relative to tumor volume at 14 weeks of age in depleted or not-depleted NeuT+ mice. **b.** Principal Component Analysis (PCA) of FACS-sorted neutrophils from NeuT− and NeuT+ mice at preneoplastic and late stages, based on the top 10% most variable genes. **c.** ssGSEA scores of published “anti-tumoral” neutrophil signatures projected onto neutrophils from preneoplastic (PAN) and late (TAN) stages. **d.** Volcano plot showing 1572 differentially expressed genes between PAN and TAN. **e.** Over-Representation Analysis (ORA) of pathways enriched in PAN and TAN. **f.** Quantification of TGF-β by ECLIA in early and late hMG, pMG, and tMG supernatants obtained after dilaceration. **g., h.** Expression levels (log2 TPM) of *Tgfbr1* (**h.**) and *Siglecf* (**i.**) in control neutrophils from NeuT− at both stages (Ctl NeuT−), in PAN and TAN. **i.** ssGSEA score of the published “SiglecF^high^ vs SiglecF^low^ neutrophil” signature (based on up- and downregulated DEGs), reflecting a tumor-promoting neutrophil population in lung tumor. Data information : (**c., i.**) Mann-Whitney test (*p<0.05, **p<0.01, ***p<0.001); (**f.**) 2-way ANOVA test (****p<0.0001; ns p ≥ 0.05); (**g., h.**) Kruskall-Wallis test. Shown is the median ± SEM. (**p<0.01, ***p<0.001; ****p<0.0001; ns p ≥ 0.05)

In the objective to better understand the role of PAN at preneoplastic stage of BC, we performed bulk RNA-sequencing (RNA-seq) of sorted PAN, along with their temporal controls from NeuT− mice and TAN (see Materials and Methods section). Using unsupervised analysis, we showed that PAN were closer to healthy neutrophils (hNeu) whereas TAN clustered together (**Fig 1b**). Building on the work of Ng et al., who functionally defined anti-tumoral T2 and pro-tumoral T3 neutrophils and generated their corresponding transcriptional signatures^9^, we examined how PAN and TAN relate to these validated states. In line with depletion results (**Fig 1a; Fig S2e**), we found that PAN were enriched in the T2 (anti-tumoral) signature (**Fig 1c**), whereas TAN displayed higher ssGSEA scores for the T3 (pro-tumoral) signature (**Fig S2f**). These findings were further supported by additional published murine neutrophil signatures^7,8^, which similarly confirmed the anti-tumoral/canonical profile of PAN (**Fig 1c**) and the pro-tumoral profile of TAN (**Fig S2f**). Then, we compared PAN and TAN and identified 822 differentially expressed genes (DEGs) upregulated in PAN and 750 in TAN (**Fig 1d**). Using Gene Ontology (GO) analysis on the DEGs, we demonstrated than PAN featured pathways of canonical neutrophil activities, including activation, phagocytosis, reactive oxygen species (ROS) production and degranulation (**Fig 1e**). Regarding TAN, they exert translation, glycolysis and protein folding as described in a recent study in pancreatic cancer^9^ (**Fig 1e**). We also observed that TAN are enriched in TGF-β signaling (**Fig 1e**). Consistently, TGF-β concentrations in the supernatant of mechanically dissociated MG were higher in tMG compared to pMG and hMG (**Fig 1f**). This enrichment is further supported by their overexpression of the TGF-β receptor gene *Tgfbr1* (**Fig 1g**) and the upregulation of the TGF-β target gene *Siglecf*^40^ (**Fig 1h**). Interestingly, TAN are enriched in pro-tumoral SiglecF^high^ neutrophils (**Fig 1i**), which express genes related to cancer-promoting processes, including T cell immunosuppression, angiogenesis, extracellular matrix remodeling, myeloid cell recruitment, and tumor proliferation, as previously defined^40–42^. Altogether, our findings demonstrate that PAN exert anti-tumoral/canonical activities at the preneoplastic stage, and shift toward a pro-tumor phenotype at late stage.

### Preneoplastic epithelial cells exhibit UPR activation and immunogenic features *in vivo*

To characterize the interaction between PAN and preneoplastic epithelial cells (pEpi), we sorted pEpi and performed bulk RNA-seq (see Materials and Methods section). By unsupervised analysis, we observed that epithelial cells from NeuT− mice (referred to as healthy epithelial cells – hEpi) at both early and late stages clustered together, whereas preneoplastic (pEpi) and tumor epithelial cells (tEpi) formed 2 distinct clusters. Tumorigenesis process explained the PC1 (53,5%), showing progression from hEpi to pEpi and ultimately to tEpi. Interestingly, PC2 (13,2%) distinguishes pEpi from others (**Fig 2a**) suggesting that pEpi exhibit a distinct transcriptional program which is a transitional state between hEpi and tEpi. This result has been validated by hierarchical clustering distinguishing transcriptomes between pEpi to hEpi and tEpi based on the 10% most differential genes (**Fig 2b**). Looking closer to gene expression, we observed different patterns across tumorigenesis. Indeed, the gene cluster A (red) showed a progressive decrease (**Fig S3a**), while the cluster C (blue) exhibited the opposite trend (**Fig S3b**). Regarding the cluster D (purple), it is already expressed at the preneoplastic stage and persisted into the late stage (**Fig S3c**), suggesting that these genes are associated with tumor initiation and maintenance. For the cluster B (green), it appears to be specific to pEpi, with a transient overexpression at this stage (**Fig 2b; Fig S3d**). Interestingly, overall-representation analysis (ORA) revealed that pEpi expressing cluster C genes (**Fig S3b**) were enriched for pathways related to cellular stress and metabolic reprogramming, reflecting their adaptation to oncogenic transformation, as illustrated by upregulated KRas signaling (**Fig 2c**). This cellular stress activates the endoplasmic reticulum (ER) adaptive response, the unfolded protein response (UPR – **Fig 2c–d**). In addition, pEpi display immunogenic features: cluster B genes are involved in inflammatory responses, IFN-γ signaling, and allograft rejection (**Fig 2e**), while cluster D exhibits an interferon response signature, indicating that an immune response is already initiated at the preneoplastic stage (**Fig S3e**). Consistently, pEpi overexpressed *Cxcl1* (**Fig S3f**), and direct quantification confirmed that CXCL1 concentrations were already higher in pMG than in hMG (**Fig S3g**). This increased CXCL1 abundance is consistent with the early tendency toward neutrophil recruitment observed at the preneoplastic stage (**Fig S1j**)

**Figure 2:**
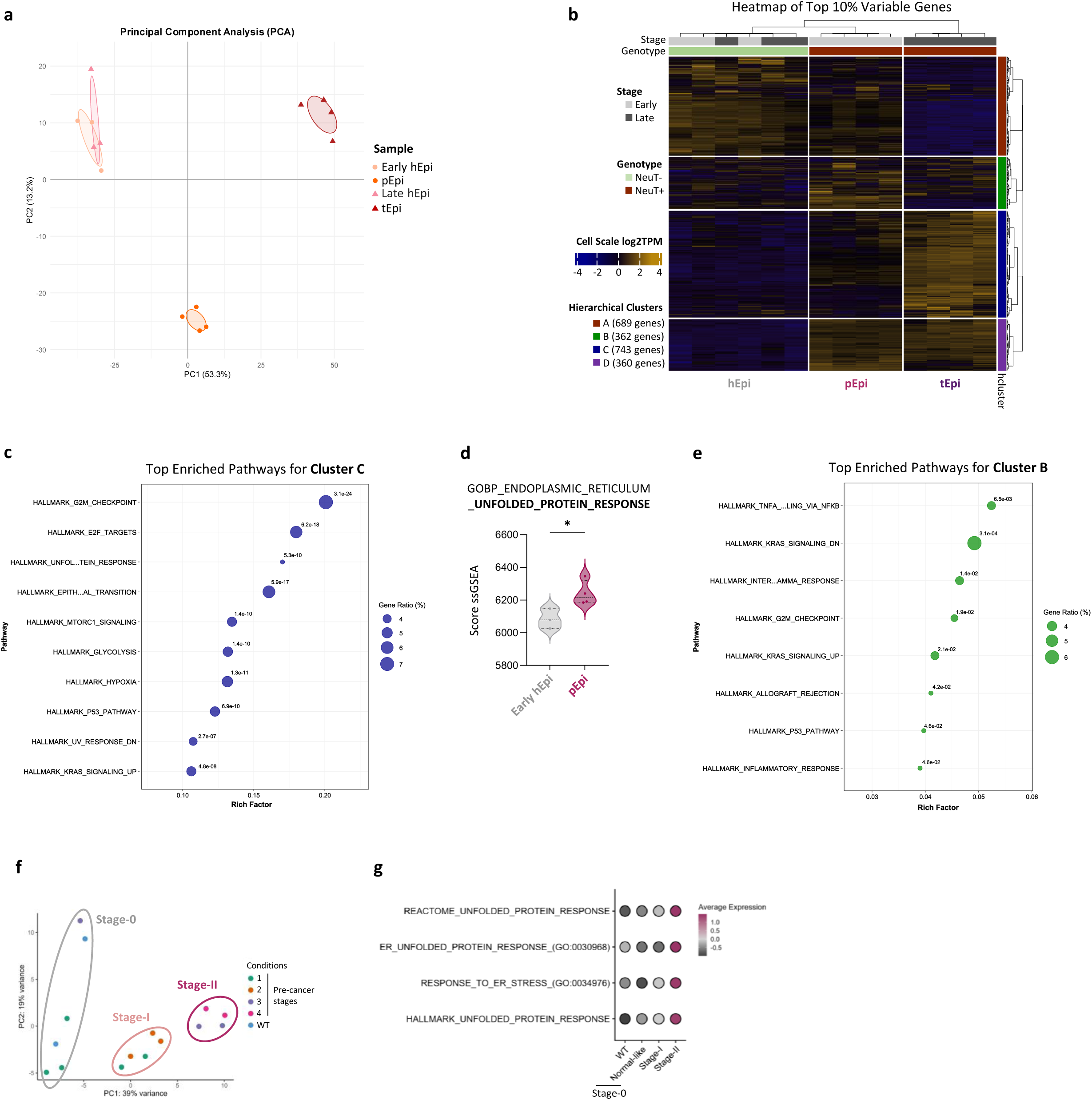
Preneoplastic epithelial cells activate UPR pathways and acquire an immunogenic phenotype. **a.** Principal Component Analysis (PCA) of FACS-sorted epithelial cells from NeuT− and NeuT+ at preneoplastic and late stages (PC1 and PC2), based on the top 10% most variable genes. **b.** Hierarchical clustering of FACS-sorted epithelial cells from NeuT− and NeuT+ mice at preeoplastic and late stages, revealing four distinct transcriptional profiles. **c.** Over-Representation Analysis (ORA) of Hallmark gene sets from the MSigDB database performed on genes from the Cluster A. **d.** Violin plot representing ssGSEA enrichment score of the UPR signature in healthy (hEpi, n=3) and preneoplastic (pEpi, n=4) epithelial cells in the MMTV-Neu mouse model. **e.** ORA of Hallmark gene sets from the MSigDB database performed on genes from the Cluster D. **f.** PCA of luminal epithelial cells from previous wild-type (WT) or pre-cancer stages annotation. Stage-0 (WT and Normal-like cells), Stage-I and Stage-II were manually annotated in BLG-Cre, BRCA1^f/f^, p53^+/−^ mouse model. **g.** Dot plot representing ssGSEA enrichment scores of ER stress and UPR response signature in luminal epithelial cells from WT, Normal-like, Stage-I, and Stage-II in the BLG-Cre, BRCA1^f/f^, p53^+/−^ mouse model. Data information: (**d.**) Welch’s t test (*p<0.05).

To validate our findings in another BC model, we extended analyses to an independent publicly available scRNA-seq dataset from a triple negative BC (TNBC) mouse model (BLG-cre; BRCA1^f/f^; p53^+/−^), which includes wild-type (WT) tissue and pre-cancerous stages^38^. We conducted pseudo-bulk analyses and performed principal component analysis (PCA), revealing three distinct groups of progenitor cells: a WT-like group, closely resembling WT progenitors, and 2 additional pre-cancerous groups, designated as stages I and II (**Fig 2f**). Interestingly, ssGSEA analysis revealed an increase of UPR signature scores starting at stage I, with a pronounced amplification at stage II (**Fig 2g**). Using the same mouse model available in our laboratory, we performed flow cytometry on preneoplastic tissue and showed neutrophils recruitment at early stage (**Fig S3h**), similar to the MMTV-*Neu* model. Altogether, these results indicate that preneoplastic epithelial cells are under cellular stress, activate the UPR, and display immunogenic features, highlighting their potential role in early neutrophil recruitment.

### Oncogenic stressed epithelial cells trigger UPR *in vitro*

To investigate the initial interactions between epithelial cells and neutrophils, we focused on the oncogenic stress (OS)-driven phase and developed an *in vitro* model using primary immortalized human mammary epithelial cells (HMEC) undergoing oncogene-induced senescence (OIS, referred to as sHMEC – **Fig S4a**) following HRas^G12V^ overexpression (doxycycline induced, **Fig S4b**) and activation of its signaling pathway (**Fig S4b-c**), which is downstream of HER2 amplification in breast cancers^43^. Following 5 days of Ras pathway activation, senescent HMEC display classical senescence features, including morphological changes (**Fig S3d**), proliferation arrest (**Fig S4e-f**), cell-cycle arrest (**Fig S4g**), β-galactosidase activity (**Fig S4h**) and transcriptional downregulation of the DNA damage response^44^ (DDR - **Fig S4i**). Using our transcriptomic data, we found that sHMEC exhibit senescence signatures from public datasets^45,46^ (**Fig S4j**) as well as genes encoding senescence-associated secretory phenotype (SASP) factors (**Fig S4k**). Interestingly, prior to senescence (**Fig S4d-j**), HMEC experience OS (**Fig S4a**), hereafter referred to as early HMEC (eHMEC), as evidenced by a significant enrichment of a stress response signature (**Fig 3a**) and expression of NKG2D ligands (NKG2D-L - ULBP2-5-6) that have been shown to be expressed by stressed cells^47^ (**Fig 3b**). Transcriptomic analyses revealed that this stress is an endoplasmic reticulum (ER) stress (**Fig 3c**), triggering the UPR, which encompasses three signaling pathways initiated by IRE1, PERK, and ATF6^48^ (**Fig S5a**). Transcriptomic analysis revealed the enrichment of IRE1- and PERK-target signatures (**Fig 3d**), validated by western blot showing IRE1 overexpression and PERK/EIF2A phosphorylation, with no change in ATF6 (**Fig 3e**). Upregulation of IRE1 (spliced form of *xbp1* (*xbp1*s), *dnajb9*, *hrd1* and *herpud* - **Fig 3f**) and PERK targets (*chop* and *ppp1r15a* – **Fig 3g**) was confirmed in eHMEC compared to control HMEC expressing inducible GFP (**Fig S5b-c**). We further showed that this UPR activation is specific to OS as (i) physiological Ras activation via EGF^49,50^ (5–100 ng/mL, 0.5–72 h) did not induce UPR target genes (**Fig S5d-f**), and that (ii) although ER stress can result from glucose deprivation^51,52^, in HMEC-HRas the culture medium was not limiting, and glucose supplementation did not affect IRE1 or PERK activation (**Fig S5g-i**). Together, these results show that eHMEC activate the UPR pathways, recapitulating the response observed in pEpi *in vivo*, making them a relevant model to study the immunogenicity of early transformed mammary epithelial cells and their interactions with neutrophils.

**Figure 3:**
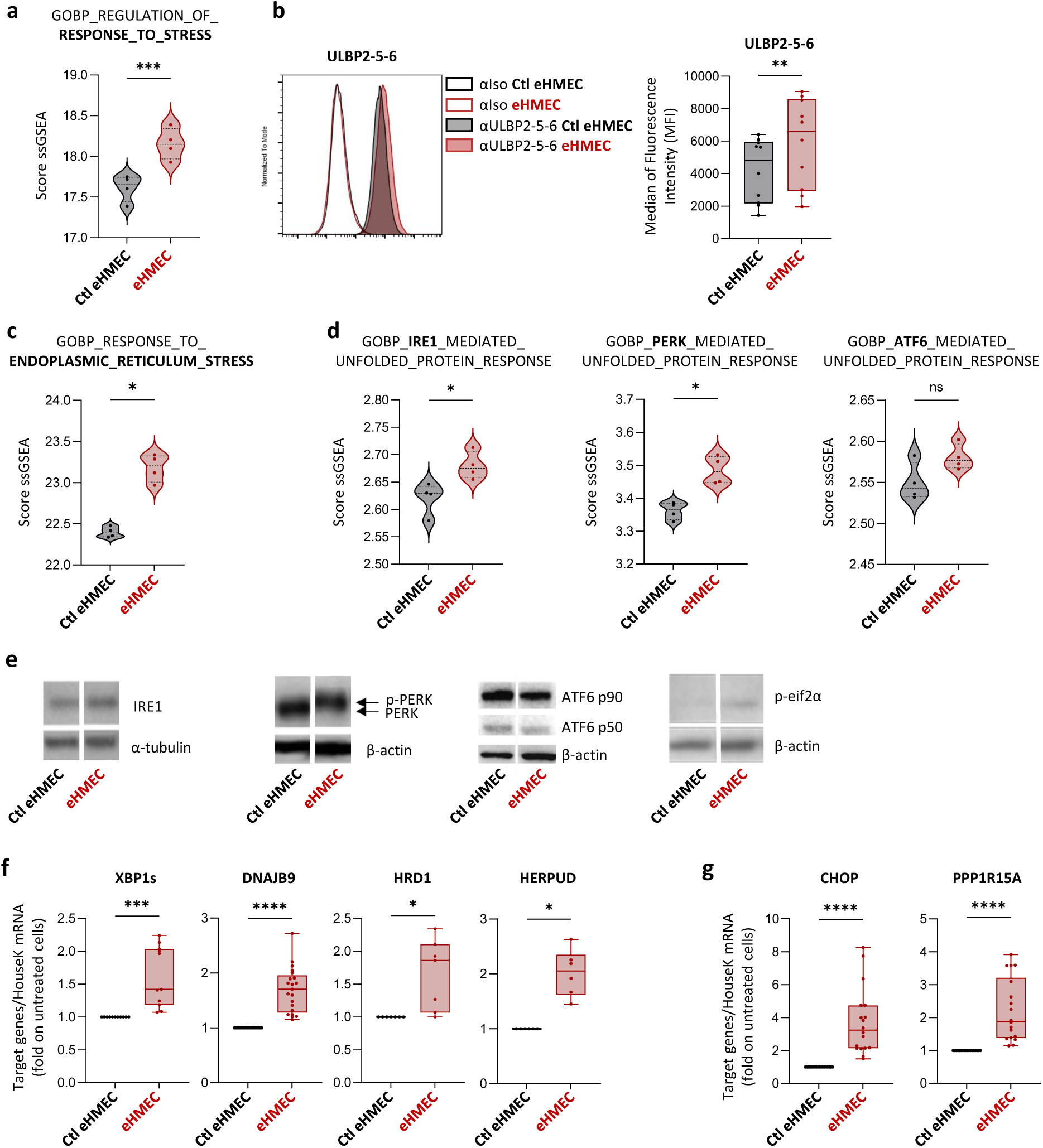
At the preneoplastic stage, HMEC are characterized by ER stress leading to UPR activation. **a.** Violin plot representing ssGSEA enrichment score of Response to Stress in Ctl eHMEC and eHMEC (n=4) using microarray data. **b.** Representative histogram plot of ULBP2-5-6 expression monitored by flow cytometry in Ctl **e**HMEC and eHMEC (left panel). Box plot illustrating the median of fluorescence intensity (MFI) of the stress ligands expression (right panel) (n=10). **c., d.** Violin plots representing ssGSEA enrichment score of the ER stress Response (**c.**) or the IRE1 (left panel), PERK (central panel) and ATF6 (right panel) pathways (**d.**) signatures in Ctl eHMEC and eHMEC (n=4) using microarray data. **e.** Representative western blot analysis of Unfolded Protein Response (UPR) sensors (IRE1, PERK and ATF6) and eiF2⍺ phosphorylation (downstream of PERK) using whole cell lysates from Ctl eHMEC and eHMEC. **f., g.** Box plots representing IRE1 (**f.**) and PERK (**g.**) target genes expression measured by RT-qPCR and normalized on Ctl eHMEC. |Data information: (**a., c.-d**) Mann-Whitney test (*p<0.05; ***p<0.001; ns p≥0.05). (**b., f.-g.**) Two-tailed Wilcoxon test. Shown is the median ± SEM (*p<0.05; ***p<0.001; ****p<0.0001).

### eHMEC exhibit immunogenic features modulated by the UPR

To pursue the parallel with the *in vivo* model (**Fig S3f**), we next investigated whether eHMEC also exhibit immunogenic features. Performing DEGs analysis on the transcriptomes of eHMEC compared to untreated HMEC (Ctl eHMEC), we identified 183 genes including 35 (19%) associated with secreted proteins, 46 (25%) related to plasma membrane proteins, 9 (5%) that are both secreted and plasma membrane-bound, and 93 (51%) related to proteins localized in other subcellular compartments (cytosol, ER, etc. – **Table S1** and **Fig 4a**, Uniprot database (https://www.uniprot.org/)). The top 10 DEGs of the secreted proteins’ group contains several immunogenic chemokines (*cxcl8/il-8*, *cxcl3*, *cxcl1* and *cxcl2*) and cytokines (*csf2*) (**Fig 4b**; **Fig S4k**) that were also upregulated at the protein level as quantified in the eHMEC supernatant by ELISA assay (**Fig 4c**). Regarding type I interferons (IFNs), which are highly immunogenic molecules^53^, we detected the enrichment of a type I-IFN signature (**Fig 4d**) and the expression of interferon-stimulated genes (ISGs; **Fig 4e**) in our transcriptomic data of eHMEC, consistent with our previous observation *in vivo* of these features in the cluster D (**Fig 2b; Fig S3e**). Among membrane ligands, in addition to NKG2D-L which are also reported for their immunogenic potential in promoting the elimination of senescent tumor cells by NK cells^54^, we identified an increase of *icam1* in the DEGs (**Table S1, Fig 4f**), confirmed at the protein level in eHMEC (**Fig 4g**). PD-L1 (*cd274*), which is also known to be regulated by the Ras pathway^55^ and expressed by senescent cells^56^, is overexpressed by eHMEC in transcriptomic data (**Table S1, Fig 4h**) as well as at the protein level by flow cytometry (**Fig 4i**). Taken together, our findings suggest that eHMEC are highly immunogenic due to their immunoactive secretome and the expression of innate immune-related membrane ligands.

**Figure 4:**
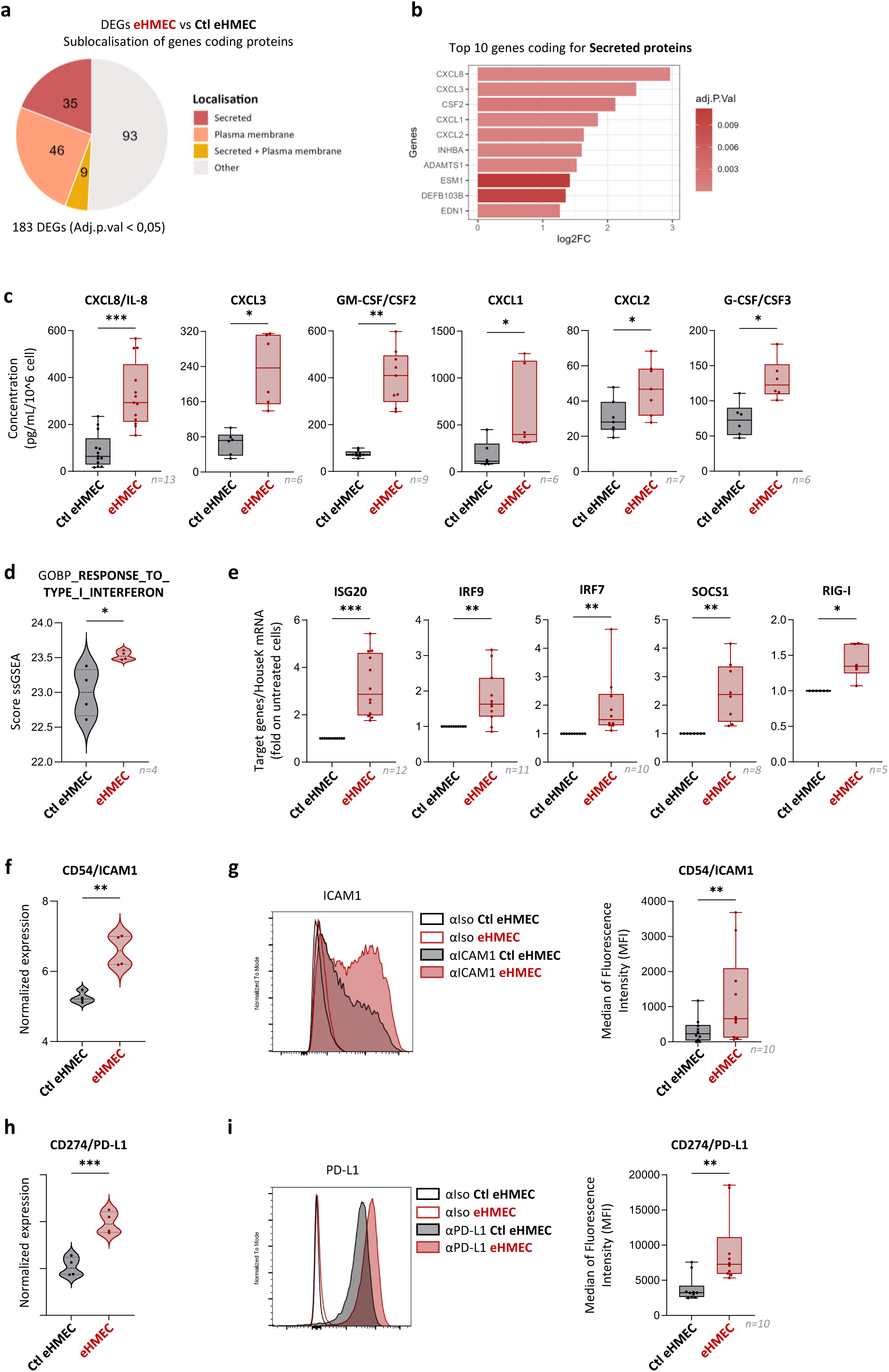
eHMEC induces the expression of immunogenic secretome and membrane ligands. **a.** Pie chart representing the subcellular localization of differential expressed protein-coding genes from the comparison of eHMEC versus Ctl eHMEC using microarray data. **b.** Top 10 of secreted protein-coding genes among DEGs between eHMEC and Ctl eHMEC. **c.** Quantification by ELISA of cytokines and chemokines production in Ctl eHMEC and eHMEC supernatants. Concentrations were normalized for 10^6^ cells (n=6 to 13). **d.** Violin plot representing ssGSEA enrichment score of the Response to type I Interferon in Ctl eHMEC and eHMEC (n=4). **e.** Interferon-stimulated genes (ISG) expression were measured by RT-qPCR and normalized on Ctl eHMEC (n=5 to 12). **f.** Violin plots representing the normalized expression of CD54/ICAM1 genes in Ctl eHMEC and eHMEC (n=4). **g.** Representative histograms and box plots showing the MFI of ICAM1 in Ctl eHMEC and eHMEC (n=10). **h.** Violin plots representing the normalized expression of CD274/PD-L1 genes in Ctl eHMEC and eHMEC (n=4). **i.** Representative histograms and box plots showing the MFI of PD-L1 in Ctl eHMEC and eHMEC (n=10) Data information: (**c., e., g., i.**) Two-tailed Wilcoxon test. Shown is the median +/− SEM (*p<0.05, **p<0.01, ***p<0.001), (**d., f., h.**) Mann-Whitney test (*p<0.05, **p<0.01, ***p<0.001)

The UPR pathways have been shown to regulate the immunogenicity of ER-stressed cells by controlling the secretion of cytokines and chemokines^57,58^, as well as the expression of immunogenic markers like NKG2D ligands^59^ and the exposure of calreticulin^60,61^. To assess the involvement of PERK and IRE1 pathways in regulating the immunogenic potential of eHMEC, we used the pharmacological inhibitors GSK-2606414 (PERK-specific) and 4µ8c (IRE1-specific) (**Fig S6a-c**). Treatment with GSK-2606414 had no impact on the secretion of neutrophil-attracting chemokines IL-8, GM-CSF, and CXCL1 by eHMEC (**Fig S6d**) whereas 4µ8c significantly inhibited their secretion (**Fig S6e**). Both inhibitors did not impact significantly the basal secretions by untreated HMEC (**Fig S6d-e**). Regarding membrane-bound immunogenic markers, ICAM1 expression is unaffected by the inhibitors whereas the basal expression of PD-L1 is decreased by the 4µ8c treatment. For ULBP2-5-6, PERK inhibition reduced their basal expression, while IRE1 inhibition amplified the eHMEC-induced expression (**Fig S6f-g**). In summary, these results show that eHMEC activate UPR pathways and that the IRE1 pathway specifically modulates neutrophil-related immunogenic features, thereby recapitulating the phenotype observed *in vivo* in pEpi.

### eHMEC immunogenicity triggers specific neutrophil recruitment and activation

We next investigated whether eHMEC are able to recruit neutrophils. Using red cell depleted whole blood from healthy donors, as a source of immune cell populations (lymphoid and myeloid cells) we performed transwell assays in presence of supernatants of eHMEC or untreated cells (**Fig 5a**). Interestingly, the only immune population significantly recruited by the eHMEC supernatant is neutrophils (**Fig 5a**). This selectivity was confirmed using peripheral blood mononuclear cells (PBMCs) (**Fig S7a**) and purified neutrophils, demonstrating a direct effect of eHMEC supernatant on their migration (**Fig 5b**). Next, we evaluated neutrophil recruitment *in vivo* using a matrigel plug assay^39^ (**Fig S7b**). We first confirmed *in vitro* that matrigel did not affect HRas activation (**Fig S7c**), HMEC proliferation (**Fig S7d**), or secretome production (**Fig S7e**). Then we engrafted at d0 matrigel-enriched with dox or not (control) and containing inducible HMEC in nude mice, and we harvested the cells at d3 (eHMEC - **Fig 5c**). Interestingly, we showed a selective increase in terms of frequency and number of neutrophils among immune cells in OS condition compared to untreated condition *in vivo* (**Fig 5d**), while the frequency and number of immune cells among viable cells remained unchanged (**Fig S7f**).

**Figure 5:**
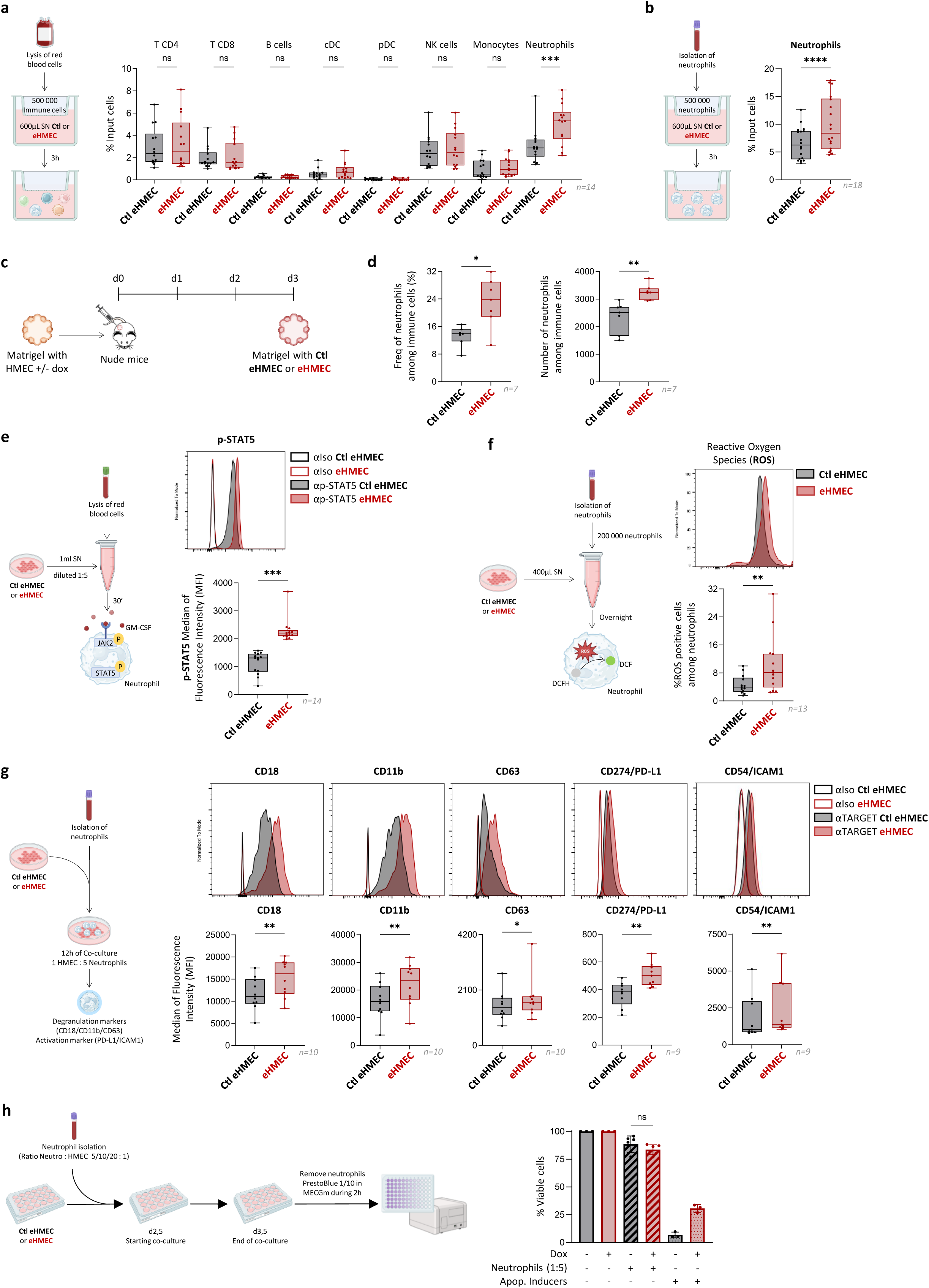
Neutrophils sense and are activated by the immunogenicity of eHMEC a., **b.** Schematic representation of transwell assays performed with total immune cells (**a.**) or isolated neutrophils (**b.**) (left panels). Box plots showing the recruitment of immune cell populations (**a.**, n=14) or neutrophils (**b.**, n=18) by Ctl eHMEC or eHMEC normalized supernatants monitored by flow cytometry (right panels). Percentage of input cells was calculated as the ratio between the number of neutrophils in the lower compartment and the initial number of neutrophils seeded (t0) in the upper compartment. **c.** Scheme of the matrigel assay *in vivo*: HMEC were embedded in matrigel without (Ctl eHMEC) or with dox (eHMEC) and then implanted into nude mice (n=7/condition). After 3 days, the matrigel implants were recovered and neutrophils were identified by flow cytometry. **d.** Box plots representing the frequency (left panel) or the number (right panel) of neutrophils among immune cells in matrigel containing Ctl eHMEC or eHMEC, assessed by flow cytometry (n=7). **e.** Schematic representation of the p-STAT5 assay using total immune cells stimulated with normalized supernatants from Ctl eHMEC or eHMEC (left panel). Box plot showing the MFI of p-STAT5 in neutrophils, as measured by flow cytometry (right panel) (n=14). **f.** Schematic representation of the reactive oxygen species (ROS) production assay using isolated neutrophils stimulated with normalized supernatants from Ctl eHMEC or eHMEC (left panel). Box plot showing the MFI of ROS in neutrophils, as measured by flow cytometry (right panel) (n=13). **g.** Schematic representation of co-culture assay performed between isolated neutrophils and Ctl eHMEC or eHMEC (ratio 5:1) (left panel). Box plot showing the MFI of degranulation (CD18, CD11b and CD63, n=10) and activation (PD-L1 and ICAM1, n=9) markers in neutrophils after co-culture, assessed by flow cytometry (right panel). **h.** Schematic representation of the 24-hour co-culture cytotoxicity assay performed between isolated neutrophils and Ctl eHMEC or eHMEC at effector-to-target (E:T) ratios of 5:1, 10:1, and 20:1 (left). Bar plot representing cell viability percentages (% viable cells) at a 5:1 ratio after treatment with an apoptosis inducer, normalized to Ctl eHMEC or eHMEC (right). Data information: (**a.-b.**, **d.-h.**) Two-tailed Wilcoxon test. Shown is the median ± SEM. (*p<0.05; **p<0.01; ***p<0.001; ****p<0.0001; ns p ≥0.05)

We next aimed to evaluate whether eHMEC activate neutrophils. Using neutrophils stimulated by eHMEC supernatant, we first showed potent STAT5 phosphorylation, notably downstream of GM-CSF receptor, with an almost 2-fold increase compared to control supernatant (**Fig 5e**), associated to a significant increase in neutrophil producing reactive oxygen species (ROS - **Fig 5f**). Last, neutrophils co-cultured with eHMEC upregulated degranulation markers including CD18, CD11b and CD63, as well as the activation markers ICAM1 and PD-L1 compared to co-cultures with control HMEC (**Fig 5g**). Then, we assessed the killing capacity of neutrophils on eHMEC by co-culture. At the effector-to-target (E:T) ratios of 5:1, 10:1, and 20:1, neutrophils did not differentially affect the viability of eHMEC versus control cells (**Fig 5h; Fig S7g-h**). Interestingly, this lack of difference in cell viability could reflect a resistance to apoptosis in eHMEC compared to control cells, as observed upon treatment with compounds known to induce apoptosis (Apoptosis Inducers) (**Fig 5h; Fig S7g-h**). Altogether, these results highlight that eHMEC specifically recruit and activate neutrophils, which remain unable to kill them in co-culture *in vitro*.

## DISCUSSION

Understanding the intrinsic and extrinsic immunosurveillance mechanisms that occur during preneoplastic stages is essential to identify critical signaling pathways that are bypassed in established tumors. These insights have a major potential to reveal novel therapeutic targets. The current study demonstrates that PAN (Preneoplastic-Associated Neutrophils) surveilled preneoplastic mammary epithelial cells characterized by the activation of the UPR and neutrophil-related immunogenic features, including an immunoactive secretome (CXCL1, IL-8, GM-CSF) and membrane ligands (ICAM1). This immunogenic profile promotes neutrophil recruitment and induces an activated state characterized by ROS production and degranulation.

PAN depletion led to reduced tumor development in approximately half of the mice (8/18) (**Fig 1a**). PAN also displayed previously reported anti-tumoral/canonical signatures and functions^7–9^ (**Fig 1c**), supporting their anti-tumoral role. The fact that approximately 50% of the animals responded is likely explained by the intrinsic asynchrony of the MMTV-*Neu* model and by inter-individual variability in the efficiency and timing of neutrophil depletion, as reflected by blood counts (**Fig S2a**). Consequently, PAN may not have been eliminated during the critical early window in all mice, which could explain the heterogeneous responses observed across individuals and the partial anti-tumor effect (**Fig 1a**). In addition, although bulk RNA-seq analyses indicate that PAN predominantly display anti-tumoral features (**Fig 1c**), neutrophils represent an heterogeneous population^7–9^, suggesting that the proportion of PAN with anti-tumoral properties may vary between mice. This highlights the potential value of scRNA-seq to better resolve neutrophil heterogeneity at preneoplastic stage in the MMTV-*Neu* model.

To better understand the sensing and the anti-tumor effect of neutrophils on tumorigenesis at preneoplastic stage, we established an innovative *in vitro* model of non-transformed human mammary epithelial cells (HMEC) undergoing OS (eHMEC), which progresses into oncogene-induced senescence (sHMEC). OIS cells are well described regarding their immunogenicity, notably via the senescence-associated secretory phenotype (SASP)^62–64^ (www.SASPAtlas.com), which can attract neutrophils capable of their clearance^65,66^. Strikingly, neutrophils sense epithelial cells as soon as they experience OS, prior to the establishment of OIS (**Fig 5a-d; Fig S7a**). This early sensing is likely orchestrated by the rapid establishment of an immunoactive secretome (**Fig 4a-c; Fig S4k**). The establishment of an OS secretome aligns with a previous study showing that stressed mouse embryonic fibroblasts (MEFs) and KRas^G12V^ hepatocytes overexpressing p21, a key regulator of the DDR^67^, release a distinct secretome preceding the SASP, referred to as the p21-activated secretory phenotype (PASP)^68^. In our study, eHMEC activate the adaptive UPR (**Fig 3c**), through the IRE1 and PERK pathways (**Fig 3d-g**) but not the DDR (**Fig S4i**). The OS secretome and, more specifically the secretion of IL-8, GM-CSF and CXCL1 are modulated in an IRE1-dependent manner (**Fig S6e**), as it has been described in MDA-MB-231 triple-negative breast cancer cell line^57^. Our findings raise intriguing questions about the preferential activation of specific adaptive intrinsic pathways, such as the UPR or DDR, depending on the cell type, the tissue origin and the driver oncogene.

In addition to secreted factors, eHMEC express immunogenic membrane ligands, including ULBP2-5-6, ICAM1, and PD-L1 (**Fig 3b**; **Fig 4f-i**). The early expression of PD-L1 is particularly intriguing, as it is classically associated with senescent cells^69,70^ and cancer cells^71^, where it promotes immune evasion by inhibiting cytotoxic T cells^56,72^ and NK cells^73^ through its interaction with PD-1. Its induction as early as the OS stage in eHMEC suggests that PD-L1 may act as an early marker of cellular stress. Moreover, the initiation of immune-escape mechanisms so rapidly during cellular transformation, particularly in Ras-driven oncogenesis, underscores the potential relevance of deploying Immune Checkpoint Inhibitors (ICIs) at preneoplastic stages to intercept tumor progression. Then, we provide further insights into PD-L1 regulation by showing that HRas-driven preneoplasia upregulates PD-L1 expression independently of UPR pathways (**Fig S6f-g**), whereas in macrophages, PD-L1 upregulation relies on IRE1 activity^74^. This UPR-independent regulation could be explained by a previous study demonstrating that activation of the Ras pathway stabilizes PD-L1 mRNA, leading to its overexpression^55^.

Interestingly, we found that neutrophil levels tend to increase at preneoplastic stage (**Fig S1j**), correlating with the upregulation of *Cxcl1* (**Fig S3f**), which has already been reported to cause neutrophil accumulation by senescent endothelial cells^75^. Then, similarly to their role at infection sites or in damaged tissues, neutrophils are the primary immune cells recruited to preneoplastic sites. Indeed, we demonstrated *in vitro* that they are the only immune cell population (**Fig 5a-b, Fig S7a**) chemoattracted by the eHMEC secretome, that includes IL-8 (**Fig 4c**) being the most potent attractor of neutrophils in humans^76^. These findings highlight neutrophils as key immune players in the detection of stressed cells and complement existing knowledge that initially emphasizes the dominant role of NK cells in this function^77^.

Once recruited upon OS, neutrophils become activated as evidenced by ROS production (**Fig 5f**) and degranulation (**Fig 5g**). While neutrophils exert anti-tumor effect *in vivo* (**Fig 1a**), they appear to be unable to eliminate eHMEC *in vitro* (**Fig 5h; Fig S7g-h**) reflecting the limitations of our simplified co-culture model. Notably, neutrophils have been shown to eliminate tumor cells through an antibody-dependent cellular cytotoxicity (ADCC) mechanism, known as trogoptosis, which relies on the Mac-1 (CD18/CD11b)/integrin axis to mediate the formation of a cytotoxic synapse^78^. This process is supported by our observation that ICAM1 is overexpressed on eHMEC (**Fig 4g**), and can interact with Mac-1 (CD11b/CD18), which is upregulated by activated neutrophils (**Fig 5g**). Then, neutrophils could recruit or activate other immune cells to kill preneoplastic cells, notably NK cells or T cells through the secretion of various chemokines, including CCL2, CCL3, and CCL4^79^. NK cells emerge as an interesting immune population, given our observation that preneoplastic cells overexpress NKG2D ligand family members (**Fig 3b**) and ICAM1 (**Fig 4g**), that are both described to drive senescent^80^ and tumor^81^ cell elimination by NK cells. Regarding T cells, neutrophils have been described as potential antigen-presenting cells, leading to T cell activation and preventing tumor growth at early stages^20,21^. Future investigations using depletion approaches could clarify the interplay between neutrophils and other immune cells and their contribution to anti-tumor immunity at preneoplastic stage. Hence, B cell depletion would impair trogoptosis by removing antibody-mediated signals; NK cell depletion would prevent neutrophil-mediated recruitment or activation of NK cytotoxicity; and T cell depletion would block neutrophil-driven antigen presentation and T cell activation, collectively impacting tumor development at early stages.

Overall, our work demonstrates for the first time that neutrophils play a key anti-tumor role in the surveillance of preneoplastic lesions. In particular, we demonstrate that neutrophils are the primary effectors of immune surveillance of preneoplastic breast cells. The activation of the UPR in these cells serves as an immunogenic warning signal detected by neutrophils. However, as tumor progress, neutrophils undergo pro-tumoral reprogramming involving, among other factors, TGF-β which ultimately contributes to tumor development. Altogether, these findings provide important insights into early BC immunosurveillance and reveal the remarkable plasticity of neutrophils as they transition from anti-tumoral to pro-tumoral states. Our work also underscores the therapeutic potential of targeting neutrophils at the preneoplastic stage. Strategies aimed at maintaining or enhancing their early anti-tumoral activity may offer a powerful means to intercept tumor initiation and prevent malignant progression.

## MATERIALS AND METHODS

### Lead contact

Further information and requests for ressources and reagents should be directed to and will be fulfilled by the lead contact, Marie-Cécile Michallet

## MATERIALS AVAILABILITY

All reagents used in this study are summarized in Table S1-S6.

All reagents generated in this study are available upon reasonable request to the lead contact.

### Data and code availability

- The microarray and RNA-seq datasets generated in this study will be deposited in GEO, and accession numbers (GSE codes) are currently pending.
- Any additional information required to reanalyze the data reported in this paper is available from the lead contact upon request

## Acknowledgments

We thank technological platforms of the CRCL, Flow cytometry core facility (CYLE), Research pathology platform East (PAR), Small animal core facility/imaging platform (P-PAC) and more specifically Thomas Barre and Sarah Urro for Imaging Procedures, Cancer genomics platform (CGP) and “Gilles Thomas” bioinformatics platform (GILLES THOMAS PLATFORM). We also thank Julien Cherfils-Vicini team at the IRCAN for its invaluable support. Our sincere thanks go to the Etablissement Français du Sang (France) for supplying the blood samples.

## Author Contributions

M.M., N.G. and L.M. developed the *in vitro* model; A.D., L.L., H.V., E.J., M.L., L.M. and M.C.M. designed the *in vitro* experiments; A.D., L.L., H.V., E.J., M.L., C.R. and L.M. performed *in vitro* experiments; J.C.V., I.M., A.D., L.M. and M.C.M. designed the matrigel experiments; J.C.V, I.M. and A.D. performed matrigel experiments; A.V. performed flow cytometry on BLG-cre; BRCA1f/f; p53+/−and analyzed scRNA-seq public data; M.P.A., P.L., L.M. and M.C.M. performed experiments on MMTV-Neu model; A.V., M.P.A., S.B. and C.A. performed *in situ* experiments and analyses; A.D., M.H., A.V., P.D. and R.P. performed bio-informatic analyses; A.D., M.P.A., L.M., C.R. and M.C.M. wrote the manuscript; A.D., M.P.A. and M.C.M. designed the figures and supplementary data; V.P., N.B.V. and C.C. provided input all along the study and on the manuscript. M.C.M. conceptualized and directed the study.

---

**Supplementary Figure 1:**
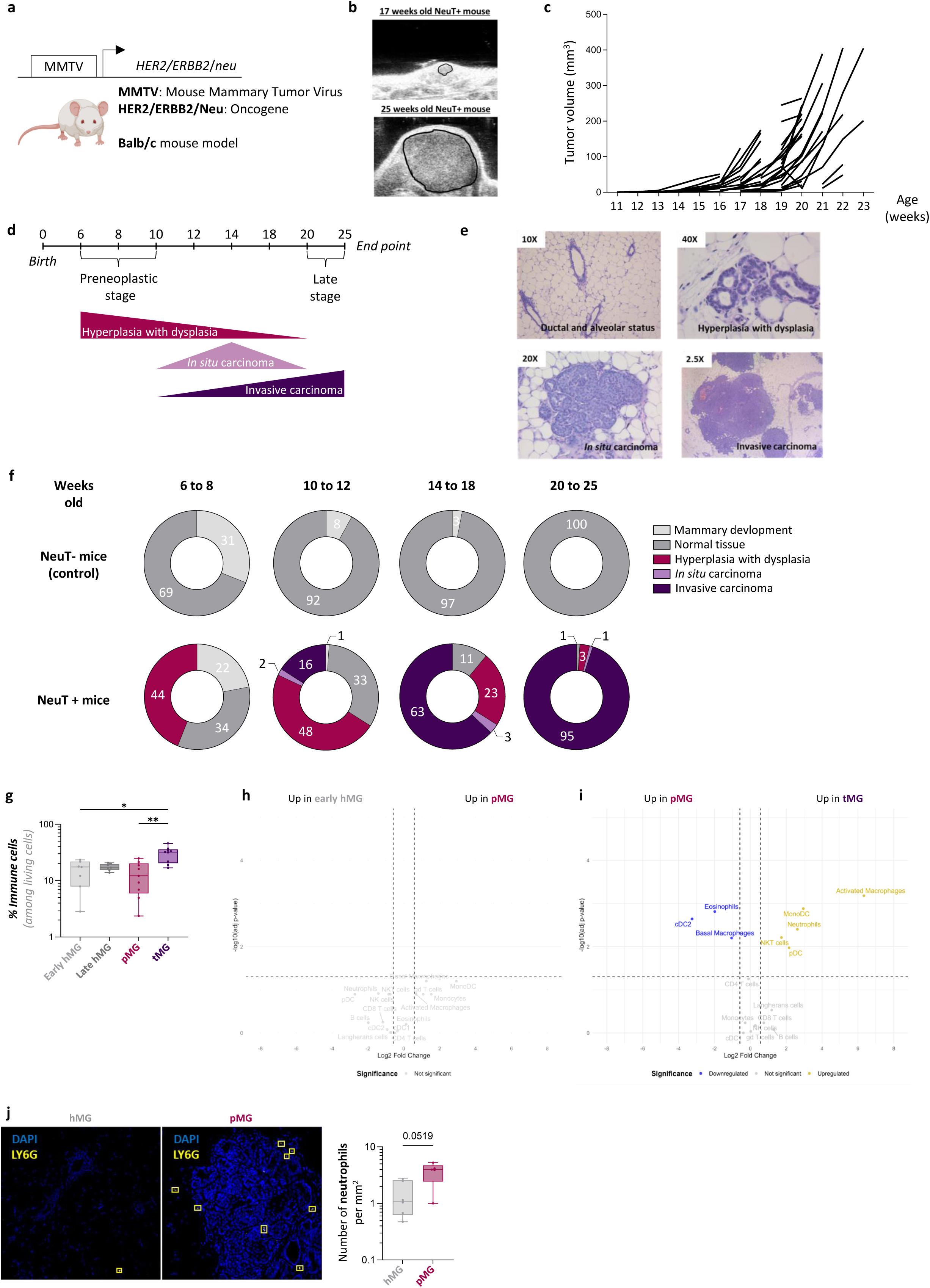
Characterization of Breast Tumor Stages and Neutrophil Infiltration in MMTV-Neu Mice. **a.** Transgenic MMTV-Neu mouse model expressing the rat HER2/neu oncogene under the transcriptional control of the MMTV-LTR promoter in a Balb/c genetic background. **b.** Ultrasound illustration of mammary gland right and left (n°1) of a NeuT+ mouse at 17 weeks old and 25 weeks old during tumor development. **c.** Ultrasound measurements of mammary glands showing the asynchrony of tumor initiation between NeuT+ mice (n=80). **d.** Timeline of mammary tumorigenesis in NeuT+ mice across weeks. Preneoplastic and late stages are determined based on mouse age, tumor volume and histological features, up to the ethical endpoint at 25 weeks. **e.** Representative Hematoxylin and Eosin staining of mammary glands from NeuT⁺ mice illustrating the successive stages of tumor development at different weeks of age. **f.** Proportion of histological features analyses of mammary glands from control (NeuT−) and tumor-bearing (NeuT+) mice across weeks (6 to 8 weeks old NeuT− (n=10) and NeuT+ (n=10); 10 to 12 weeks old NeuT− (n=12) and NeuT+ (n=14), 14 to 18 weeks old NeuT− (n=9) and NeuT+ (n=10), 20 to 25 weeks old NeuT− (n=10) and NeuT+ (n=10)). **g.** Box plots showing the proportions of immune cell among living cells in early and late healthy mammary glands (hMG), preneoplastic MG (pMG), and tumor MG (tMG). **h., i.** Volcano plots showing differential abundance of immune cell populations among immune cells between early hMG and pMG (**h.**), or between tMG and pMG (**i.**). **j.** Representative multiplex immunofluorescence staining of neutrophils (Ly6G+, yellow) in healthy mammary glands (hMG), preneoplastic MG (pMG) and tumor MG (tMG). Nuclei were stained with DAPI (blue). Quantification (cells/mm2) of neutrophils (Ly6G+) in FFPE hMG, pMG and tMG. Data information: (**g., j.**) Kruskal-Wallis test. Shown is the median ± SEM (*p<0.05, **p<0.01); (**p<0.01, ****p<0.001; ns p ≥ 0.05) (**h., i.**) Wilcoxon test.

**Supplementary Figure 2:**
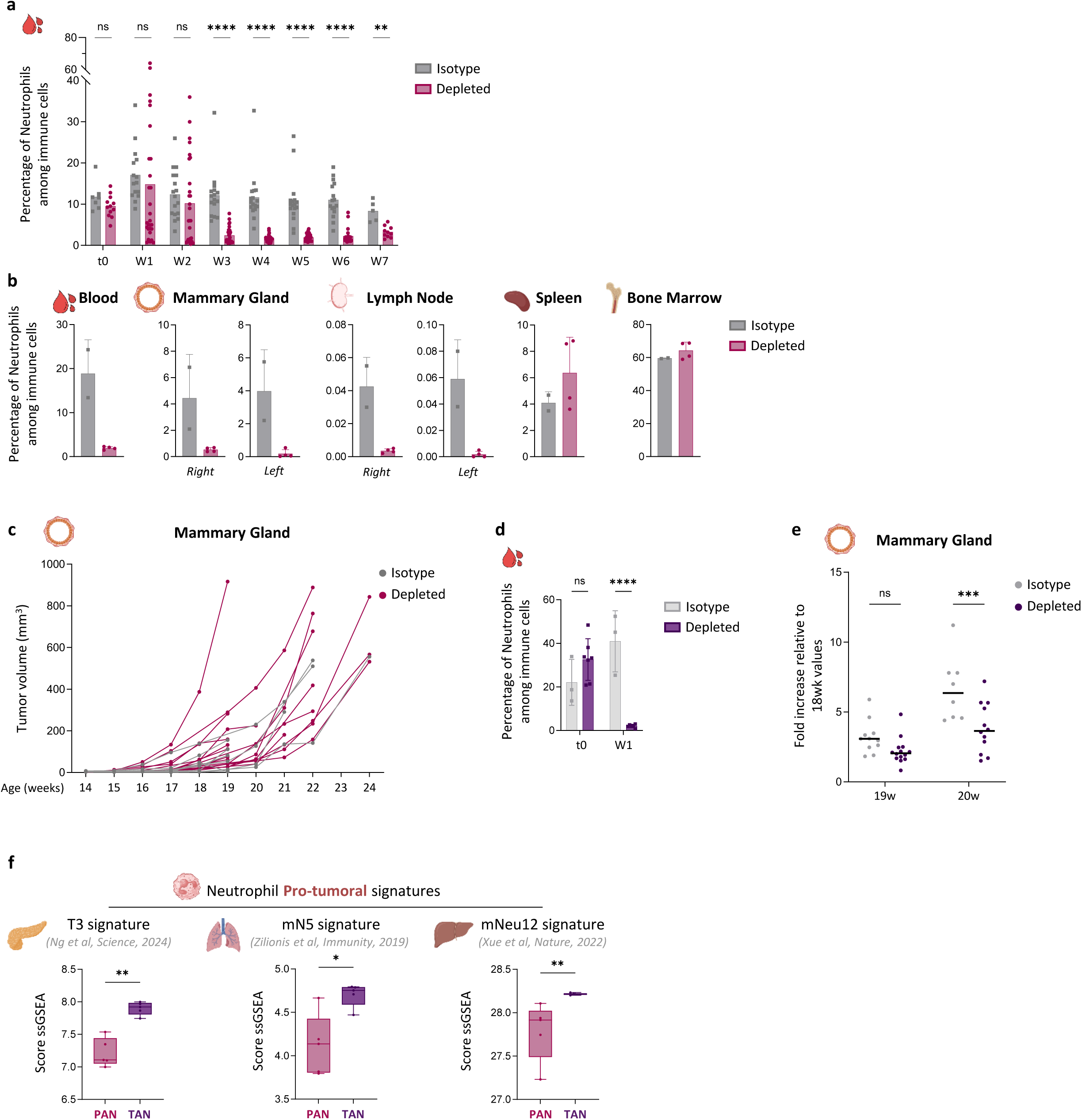
Effective depletion at early and late stages of breast cancer tumorigenesis. **a.** Proportion of neutrophils (CD11b^+^ Ly6C^int^) among viable immune cells measured at each week of depletion experiment at the preneoplastic stage (t0 at 4 to 6 weeks old, before depletion). **b.** Proportion of neutrophils (CD11b^+^ Ly6C^int^) among viable immune cells at 10-12 weeks old in blood, mammary gland (n°1, right), lymph node (left/right), spleen or bone marrow of depleted (n=4) or not-depleted (n=2) NeuT+ mice. **c.** Tumor measurement (mm^3^) by ultrasound imaging of mammary gland n°1 in control (Isotype, n =9) and neutrophil-depleted NeuT+ mice (n=18) across weeks of development after depletion at preneoplastic stage. **d.** Proportion of neutrophils (CD11b^+^ Ly6C^int^) among viable immune cells after one week of depletion at the late stage (t0 at 15 weeks old, before depletion). **e.** Monitoring of tumor growth across weeks of development during neutrophil depletion at the late stage, expressed as fold increase relative to tumor volume at 18 weeks of age in depleted or not-depleted NeuT+ mice. **f.** ssGSEA scores of published “pro-tumoral” neutrophil signatures projected onto neutrophils from preneoplastic (PAN) and late (TAN) stages. Data information: (a., d., e., f.) Mann-Whitney test (**p<0.01, ****p<0.001; ns p ≥ 0.05)

**Supplementary Figure 3:**
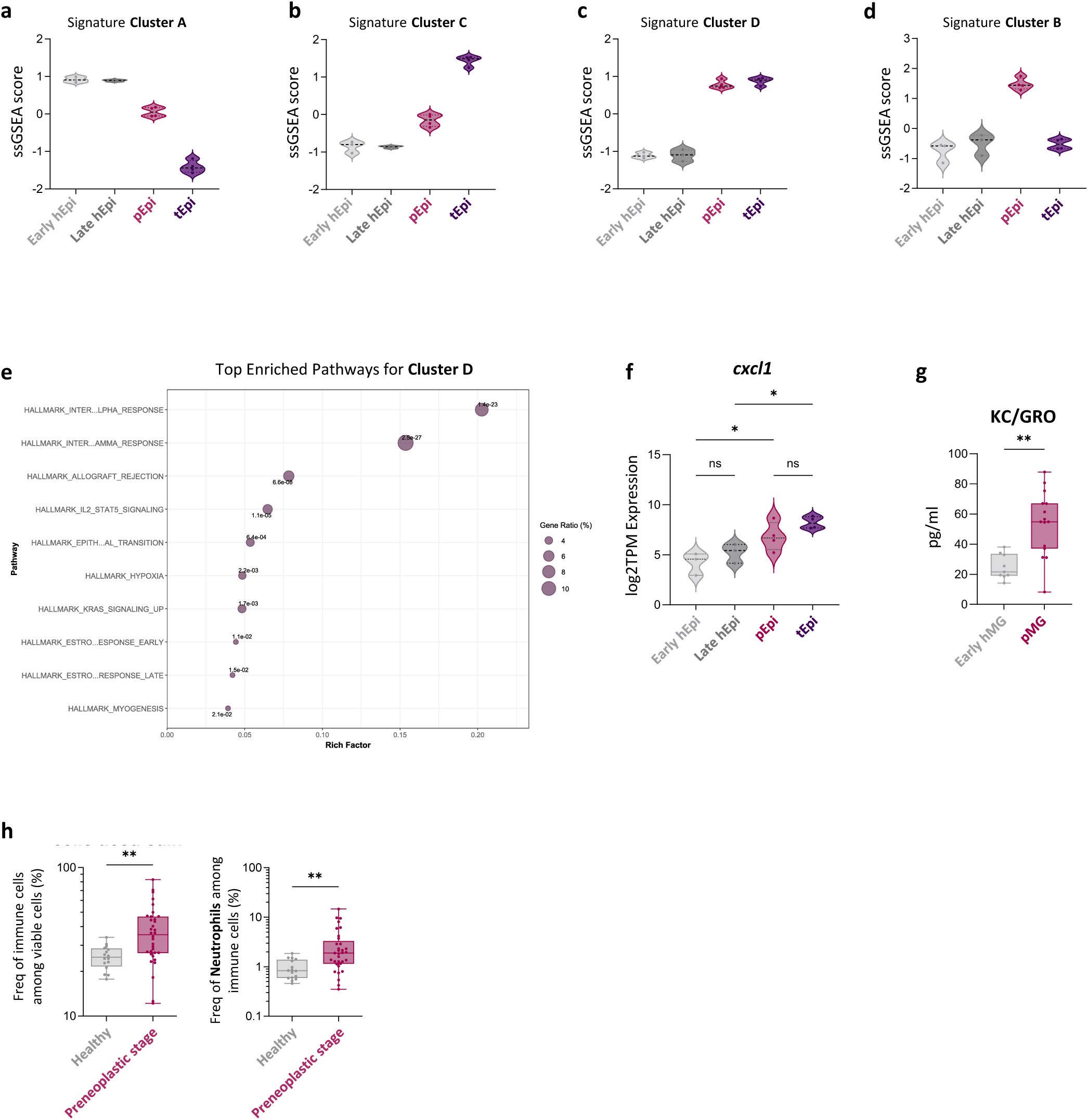
Preneoplastic epithelial cell immunogenicity promotes neutrophil recruitment. **a.**, **b.**, **c.**, **d.** Box plots representing ssGSEA enrichment score of the cluster A (**a.**), B (**b.**), C (**c.**) and D (**d.**) signatures in UPR signature early healthy (n=3), preneoplastic (n=4), late healthy (n=3) and late (n=4) epithelial cells in the MMTV-Neu mouse model. **e.** Over-Representation Analysis (ORA) of Hallmark gene sets from the MSigDB database performed on genes from the purple cluster (Cluster C). **f.** Violin plot representing the expression of *cxcl1* in early healthy (n=3), preneoplastic (n=4), late healthy (n=3) and late (n=4) epithelial cells in the MMTV-Neu mouse model. **g.** Quantification of KC/GRO by ECLIA in early hMG and pMG supernatants obtained after dilaceration. **h.** Box plot of neutrophil proportion among immune cells in healthy (n=16) and preneoplastic stage (n=34) tissues in BLG-Cre, BRCA1^f/f^, p53^+/−^ mouse model assessed by flow cytometry. Data information: (**f.**) Kruskall-Wallis test. Shown is the median ± SEM (*p<0.05, ns p ≥ 0.05) (**g.**) Mann-Whitney test (**p<0.01); (**h.**) Two-tailed Wilcoxon test. Shown is the median ± SEM. (**p<0.01)

**Supplementary Figure 4:**
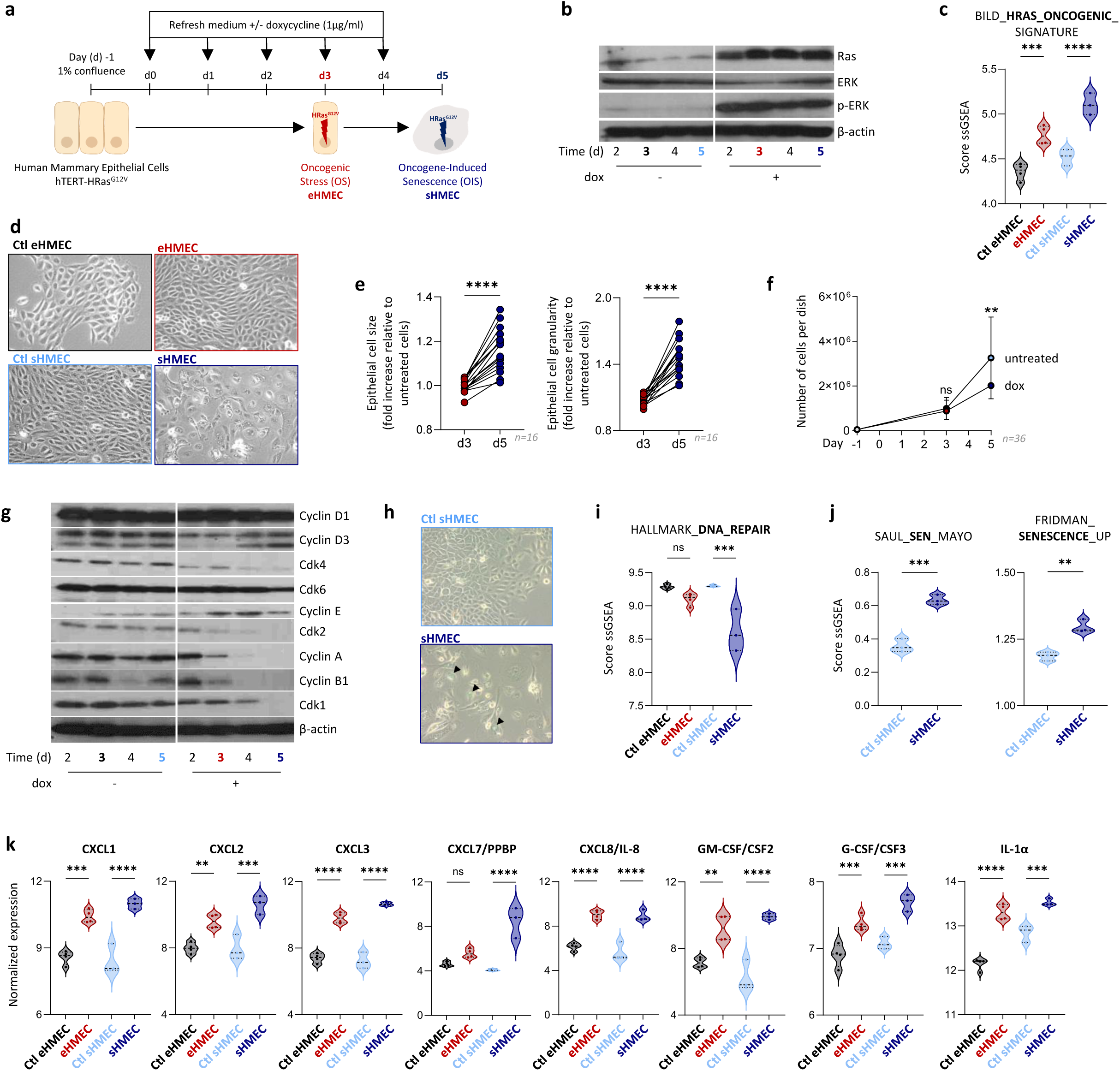
Characterization of the preneoplastic stage preceding OIS in the HMEC Model. **a.** Scheme of the kinetic induction of oncogenic stress (OS - eHMEC) and oncogene-induced senescence (OIS - sHMEC) in HMEC-HRas model. Upon doxycycline-inducible HRas^G12V^ expression, there is the preneoplastic stage at day 3 (d3 - eHMEC) followed by the senescence phase at d5 (sHMEC). **b.** Representative western blot analysis of Ras and its downstream factor ERK using whole cell lysates from a time-course experiment in untreated and doxycycline-treated HMEC-HRas. **c.** Violin plots representing ssGSEA enrichment score of the HRas Oncogenic in Ctl eHMEC and eHMEC (n=4) or Ctl sHMEC and sHMEC (n=3) using microarray data. **d.** Morphological changes in doxycycline-treated or untreated HMEC-HRas at d3 (Ctl eHMEC and eHMEC) and d5 (Ctl sHMEC and sHMEC - Magnification x20). **e.** Quantification of cell size (FSC-A; left panel) and granularity (SSCA-A; right panel) by flow cytometry of dox-treated HMEC-HRas relative to untreated cells at d3 and d5 (n=16). **f.** Mean cell number per culture dish of Ctl eHMEC, Ctl sHMEC (untreated cells), eHMEC and sHMEC (dox-treated cells - n=36). **g.** Representative western blot analysis of proteins involved in the cell cycle using whole cell lysates from a time-course experiment in untreated and dox-treated HMEC-HRas. **h.** Senescence-associated beta-galactosidase staining showing by blue coloration (arrow) in Ctl sHMEC and sHMEC. **i.** Violin plot representing ssGSEA enrichment score of the Hallmark DNA repair signature in Ctl eHMEC and eHMEC (n=4) or Ctl sHMEC and sHMEC (n=3) using microarray data. **j.** Violin plot representing ssGSEA enrichment score of a published senescence signature in Ctl sHMEC and sHMEC (n=3) using microarray data. **k.** Violin plots representing the normalized expression of cytokines and chemokines Ctl eHMEC and eHMEC (n=4) or Ctl sHMEC and sHMEC (n=3) using microarray data. Data information: (**a., j., k.**) One-Way ANOVA test (*p<0.05, **p<0.01, ***p<0.001, ***p<0.0001); (**e.**) Two-tailed Wilcoxon test. Shown is the median ± SEM (*p<0.05; **p<0.01; ***p<0.001; ****p<0.0001); (**f.**) Two-Way ANOVA test (**p<0.01; ns p≥ 0.05); (**j.**) Mann-Whitney test (**p<0.01; ***p<0.001)

**Supplementary Figure 5:**
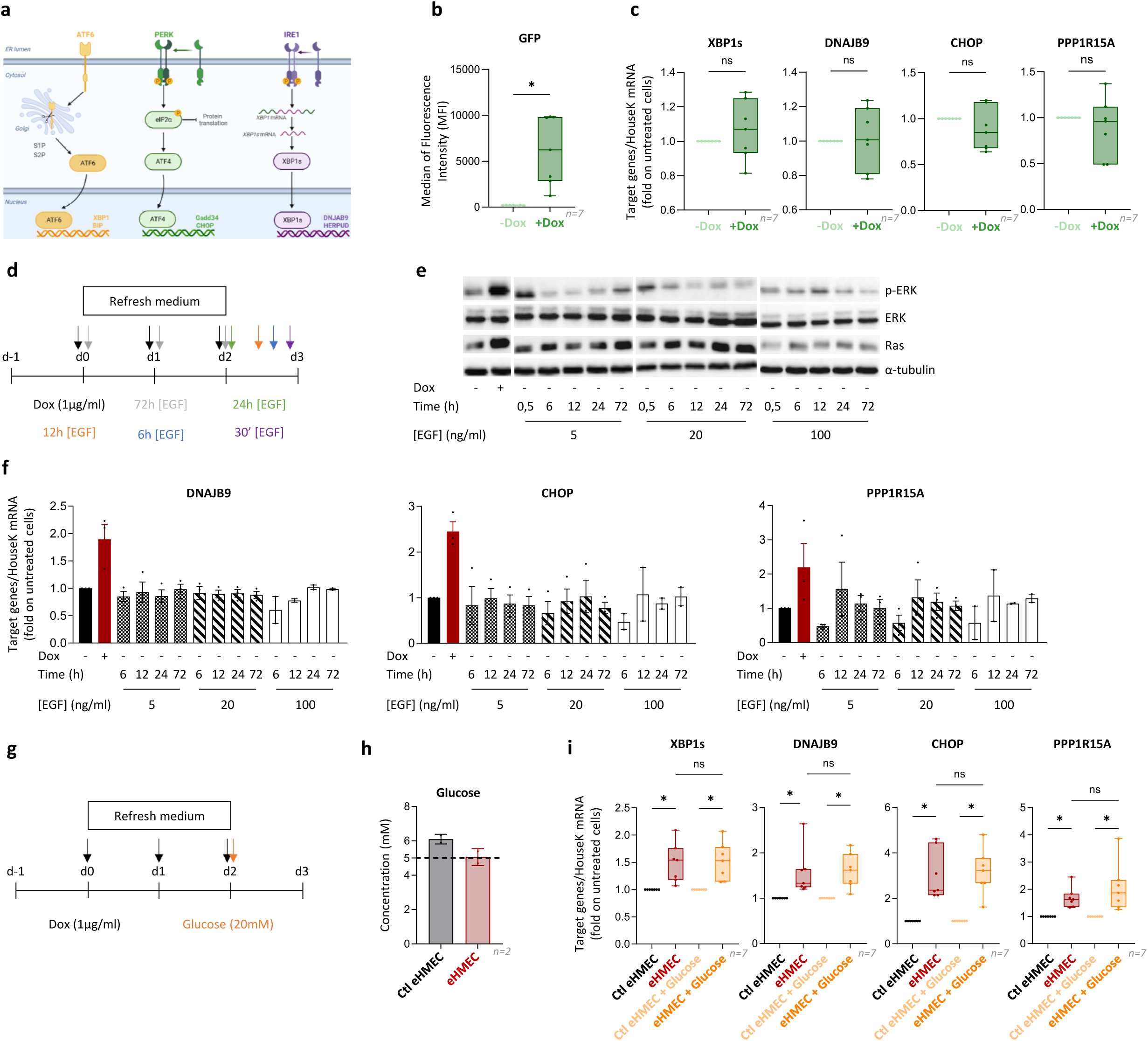
UPR pathways are specifically activated by the OS experienced by eHMEC. **a.** Scheme of the Unfolded Protein Response (UPR) pathways. **b.**, **c.** HMEC-GFP model was developed as a control for HRas overexpression. Box plot showing the MFI of the GFP expression in untreated (-Dox) or dox-treated (+Dox) HMEC-GFP at d3, assessed by flow cytometry (b.). Box plots representing IRE1 target genes (*xbp1s* and *dnajb9*) and PERK target genes (*chop* and *ppp1r15a*) expression measured by RT-qPCR and normalized on untreated HMEC-GFP at d3 (c.) (n=7). **d.** Scheme representing the protocol performed to investigate whether physiological activation of the Ras signaling pathway could elicit the same intrinsic response. HMEC-HRas cultures were supplemented with EGF at different concentrations (5, 20, and 100 ng/mL) and analyzed over time (0.5, 6, 12, 24, and 72h). For the 72-hour time point, the medium was refreshed every 24 hours with EGF at the respective concentrations. **e.** Representative western blot analysis of Ras and its downstream factor ERK using whole cell lysates from a time-course experiment in untreated, dox-treated and EGF-treated HMEC-HRas at d3. **f.** Box plots representing IRE1 target genes (*dnajb9*) and PERK target genes (*chop* and *ppp1r15a*) expression measured by RT-qPCR and normalized on Ctl eHMEC (n=3). **g.** Scheme representing the protocol performed, given that glucose deprivation is another factor capable of activating the UPR pathways. HMEC-HRas cultures were supplemented with exogenous glucose (20 mM) on d2. **h.** Bar plot representing the quantification of the glucose concentration in Ctl and eHMEC supernatants (n=2). The dashed line represents 5mM of glucose, representing a critical concentration that can induce ER stress. **i.** Box plots representing IRE1 target genes (*dnajb9* and *xbp1s*) and PERK target genes (*chop* and *ppp1r15a*) expression measured by RT-qPCR and normalized on Ctl eHMEC in presence or not of glucose, respectively (n=7). Data information: (**a.-b.**) Two-tailed Wilcoxon test. Shown is the median ± SEM (* p<0.05; ns p ≥ 0.05) (**i.**) Friedman test (* p<0.05; ns p ≥ 0.05).

**Supplementary Figure 6:**
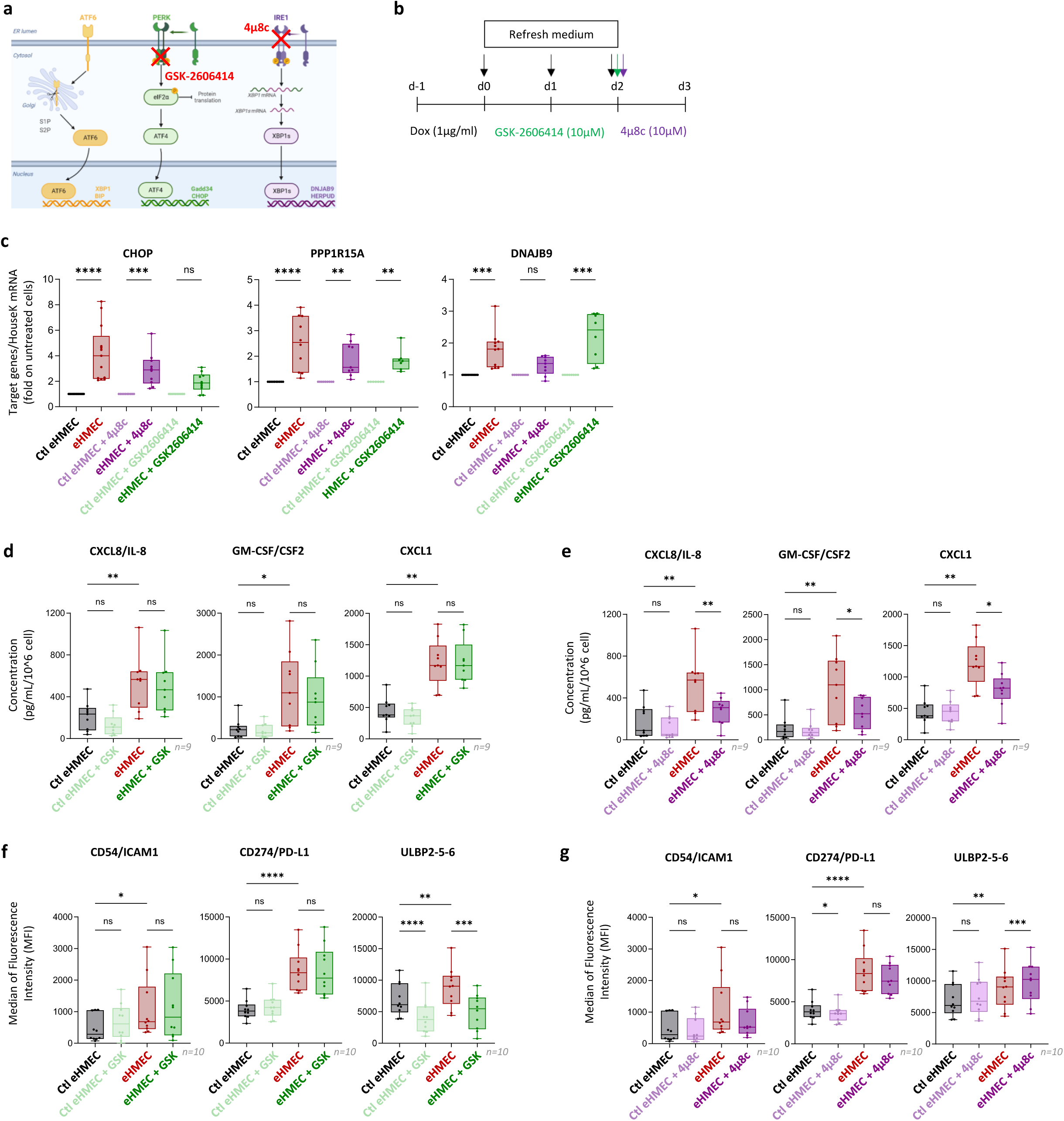
Inhibition of PERK and IRE1 pathways affect the eHMEC-induced immunogenicity. **a** Schematic illustration of the unfolded protein response (UPR) signaling pathways and their inhibition by specific inhibitors targeting the PERK and IRE1 branches. **b** Protocol for pharmacological inhibition of the PERK (GSK-2606414) and IRE1 (4µ8c) pathways. **c** Box plots representing IRE1 target genes (*dnajb9*) and PERK target genes (*chop* and *ppp1r15a*) expression measured by RT-qPCR and normalized on Ctl eHMEC in presence or not of pharmacological inhibitors (GSK-2606414 and 4µ8c), respectively. **d**, **e.** Quantification by ELISA of IL-8, CXCL1, and GM-CSF production in Ctl eHMEC and eHMEC supernatants with or without PERK inhibition (**d.**) or IRE1 inhibition (**e.**). Concentrations (pg/mL) were normalized by 10^6^ cells (n=9). **f.**, **g.** Expression of ICAM1, ULBP2-5-6, and PD-L1 in Ctl eHMEC and eHMEC with or without PERK inhibition (**f.**) or IRE1 inhibition (**g.**) assessed by flow cytometry (n=10). Data information: (**c.**) Kruskall-Wallis test. Shown is the median ± SEM. (**p<0.01, ***p<0.001; ****p<0.0001; ns p ≥ 0.05), (**d.-g.**) RM one-Way ANOVA test. Shown is the mean ± SEM. (*p<0.05, (**p<0.01, ***p<0.001; ****p<0.0001; ns p ≥ 0.05)

**Supplementary Figure 7:**
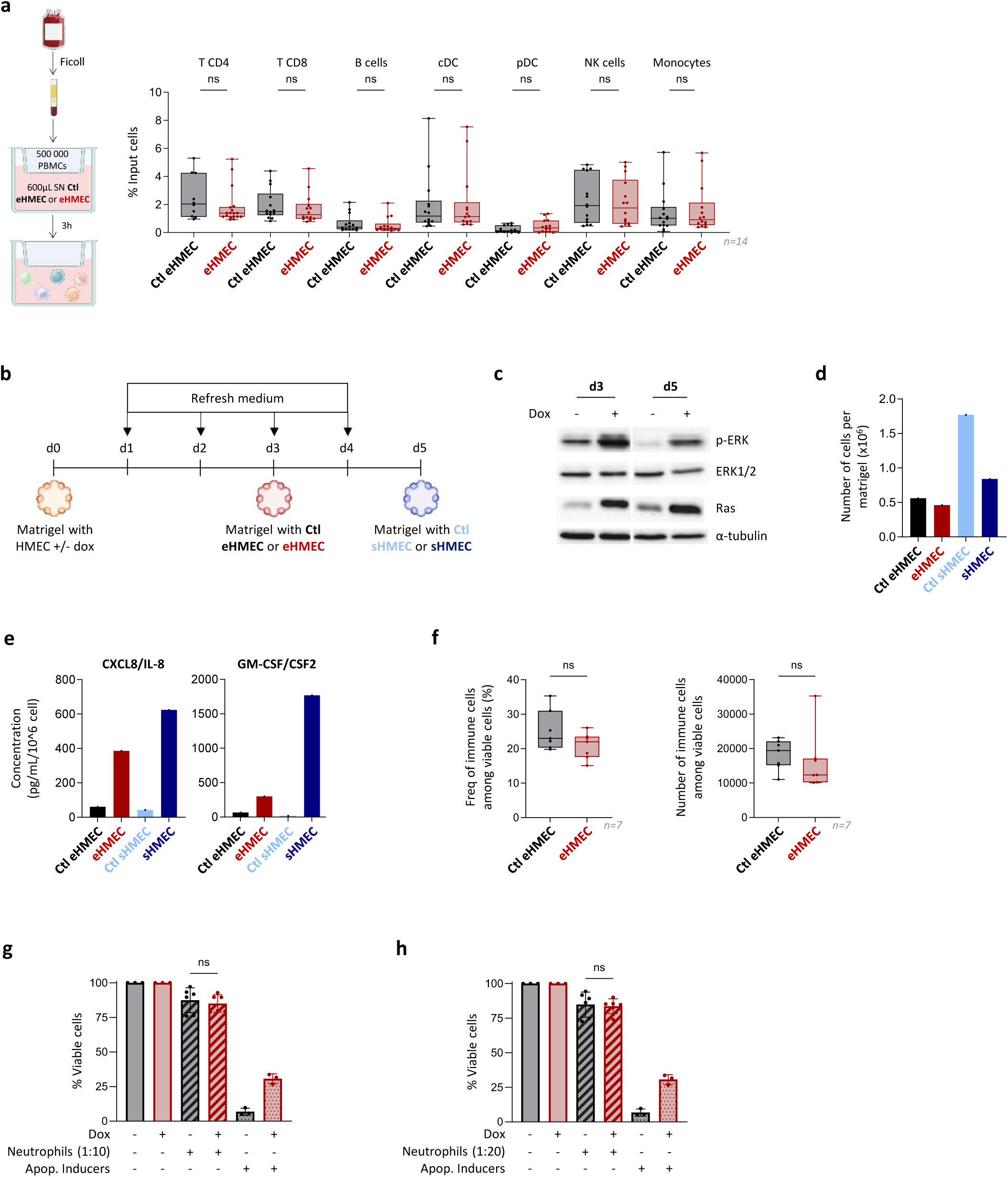
eHMEC secretome is a strong neutrophil chemoattractant. **a.** Schematic representation of transwell assays performed with using peripheral blood mononuclear cells (PBMCs) (left panel). Box plot showing the recruitment of PBMCs populations by Ctl or eHMEC normalized supernatants monitored by flow cytometry (right panel) (n=14). **b.** Scheme of the matrigel assay in vitro: HMEC were embedded in matrigel without or with dox in the culture medium and then cultured *in vitro*. Upon HRas^G12V^ induction, the OS (d3 - eHMEC) and the OIS (d5 - sHMEC) stages are determined. **c.** Representative western blot analysis of Ras and its downstream factor ERK using whole cell lysates from a time-course experiment in untreated and treated matrigel at d3 and d5. **d.** Bar plot representing the cell number per matrigel containing Ctl eHMEC, eHMEC, Ctl sHMEC and sHMEC (n=1). **e.** Bar plot representing the quantification by ELISA of IL-8 and GM-CSF production in matrigel containing Ctl eHMEC, eHMEC, Ctl sHMEC and sHMEC. Concentrations were normalized for 10^6^ cells (n=1). **f.** Box plots representing the frequency (left panel) or the number (right panel) of immune cells among viable cells in matrigel containing Ctl eHMEC or eHMEC, assessed by flow cytometry (n=7). **g.**, **h.** Bar plot representing cell viability percentages (% viable cells) at a 10:1 (**g.**) and 20:1 (**h.**) ratio, normalized to Ctl eHMEC or eHMEC without neutrophils. Data information: (**a.**, **f.-h.**) Two-tailed Wilcoxon test. Shown is the median ± SEM. (ns p≥0.05).

## REFERENCES

1. Łukasiewicz, S., et al. Breast Cancer—Epidemiology, Risk Factors, Classification, Prognostic Markers, and Current Treatment Strategies—An Updated Review. Cancers 13, 4287 (2021).

2. Sgroi, D. C. Preinvasive Breast Cancer. Annu. Rev. Pathol. 5, 193–221 (2010).

3. Chen, M.-T. et al. Comparison of patterns and prognosis among distant metastatic breast cancer patients by age groups: a SEER population-based analysis. Sci. Rep. 7, 9254 (2017).

4. Khoury, T. Preneoplastic Low-Risk Mammary Ductal Lesions (Atypical Ductal Hyperplasia and Ductal Carcinoma In Situ Spectrum): Current Status and Future Directions. Cancers 14, 507 (2022).

5. Sun, Y.-S. et al. Risk Factors and Preventions of Breast Cancer. Int. J. Biol. Sci. 13, 1387–1397 (2017).

6. Otterlei Fjørtoft, M., Huse, K. & Rye, I. H. The Tumor Immune Microenvironment in Breast Cancer Progression. Acta Oncol. 63, 33008 (2024).

7. Zilionis, R. et al. Single cell transcriptomics of human and mouse lung cancers reveals conserved myeloid populations across individuals and species. Immunity 50, 1317–1334.e10 (2019).

8. Xue, R. et al. Liver tumour immune microenvironment subtypes and neutrophil heterogeneity. Nature 612, 141–147 (2022).

9. Ng, M. S. F. et al. Deterministic reprogramming of neutrophils within tumors. Science 383, eadf6493 (2024).

10. Wang, T. et al. Tumour-activated neutrophils in gastric cancer foster immune suppression and disease progression through GM-CSF-PD-L1 pathway. Gut 66, 1900–1911 (2017).

11. Cheng, Y. et al. Cancer-associated fibroblasts induce PDL1+ neutrophils through the IL6-STAT3 pathway that foster immune suppression in hepatocellular carcinoma. Cell Death Dis. 9, 422 (2018).

12. Teijeira, Á. et al. CXCR1 and CXCR2 Chemokine Receptor Agonists Produced by Tumors Induce Neutrophil Extracellular Traps that Interfere with Immune Cytotoxicity. Immunity 52, 856–871.e8 (2020).

13. Kaltenmeier, C. et al. Neutrophil Extracellular Traps Promote T Cell Exhaustion in the Tumor Microenvironment. Front. Immunol. 12, (2021).

14. Houghton, A. M. et al. Neutrophil Elastase-Mediated Degradation of IRS-1 Accelerates Lung Tumor Growth. Nat. Med. 16, 219–223 (2010).

15. Park, J. et al. Cancer cells induce metastasis-supporting neutrophil extracellular DNA traps. Sci. Transl. Med. 8, 361ra138 (2016).

16. Xiao, Y. et al. Cathepsin C promotes breast cancer lung metastasis by modulating neutrophil infiltration and neutrophil extracellular trap formation. Cancer Cell 39, 423–437.e7 (2021).

17. Jiang, Z.-Z. et al. Neutrophil extracellular traps induce tumor metastasis through dual effects on cancer and endothelial cells. Oncoimmunology 11, 2052418.

18. Yang, S. et al. Neutrophil extracellular traps promote angiogenesis in gastric cancer. Cell Commun. Signal. CCS 21, 176 (2023).

19. Blaisdell, A. et al. Neutrophils Oppose Uterine Epithelial Carcinogenesis via Debridement of Hypoxic Tumor Cells. Cancer Cell 28, 785–799 (2015).

20. Eruslanov, E. B. et al. Tumor-associated neutrophils stimulate T cell responses in early-stage human lung cancer. J. Clin. Invest. 124, 5466–5480 (2014).

21. Singhal, S. et al. Origin and Role of a Subset of Tumor-Associated Neutrophils with Antigen Presenting Cell Features (Hybrid TANs) in Early-Stage Human Lung Cancer. Cancer Cell 30, 120–135 (2016).

22. Fridlender, Z. G. et al. Polarization of Tumor-Associated Neutrophil (TAN) Phenotype by TGF-β: “N1” versus “N2” TAN. Cancer Cell 16, 183–194 (2009).

23. Calogero, R. A., Cordero, F., Forni, G. & Cavallo, F. Inflammation and breast cancer. Inflammatory component of mammary carcinogenesis in ErbB2 transgenic mice. Breast Cancer Res. BCR 9, 211 (2007).

24. Guey, B. et al. Inflammasome Deletion Promotes Anti-tumor NK Cell Function in an IL-1/IL-18 Independent Way in Murine Invasive Breast Cancer. Front. Oncol. 10, 1683 (2020).

25. Boggio, K. et al. Interleukin 12–mediated Prevention of Spontaneous Mammary Adenocarcinomas in Two Lines of Her-2/neu Transgenic Mice. J. Exp. Med. 188, 589–596 (1998).

26. McCarthy, A. et al. A mouse model of basal-like breast carcinoma with metaplastic elements. J. Pathol. 211, 389–398 (2007).

27. Boivin, G. et al. Durable and controlled depletion of neutrophils in mice. Nat. Commun. 11, 2762 (2020).

28. Dobin, A. et al. STAR: ultrafast universal RNA-seq aligner. Bioinformatics 29, 15–21 (2013).

29. Wang, L., Wang, S. & Li, W. RSeQC: quality control of RNA-seq experiments. Bioinformatics 28, 2184–2185 (2012).

30. Patro, R., Duggal, G., Love, M. I., Irizarry, R. A. & Kingsford, C. Salmon: fast and bias-aware quantification of transcript expression using dual-phase inference. Nat. Methods 14, 417–419 (2017).

31. Gu, Z., Eils, R. & Schlesner, M. Complex heatmaps reveal patterns and correlations in multidimensional genomic data. Bioinformatics 32, 2847–2849 (2016).

32. Subramanian, A. et al. Gene set enrichment analysis: A knowledge-based approach for interpreting genome-wide expression profiles. Proc. Natl. Acad. Sci. 102, 15545–15550 (2005).

33. Castanza, A. S. et al. Extending support for mouse data in the Molecular Signatures Database (MSigDB). Nat. Methods 20, 1619–1620 (2023).

34. Hänzelmann, S., Castelo, R. & Guinney, J. GSVA: gene set variation analysis for microarray and RNA-Seq data. BMC Bioinformatics 14, 7 (2013).

35. Love, M. I., Huber, W. & Anders, S. Moderated estimation of fold change and dispersion for RNA-seq data with DESeq2. Genome Biol. 15, 550 (2014).

36. Zhu, A., Ibrahim, J. G. & Love, M. I. Heavy-tailed prior distributions for sequence count data: removing the noise and preserving large differences. Bioinformatics 35, 2084–2092 (2019).

37. Wickham, H. Data Analysis. in ggplot2 189–201 (Springer International Publishing, Cham, 2016). doi:10.1007/978-3-319-24277-4_9.

38. Bach, K. et al. Time-resolved single-cell analysis of Brca1 associated mammary tumourigenesis reveals aberrant differentiation of luminal progenitors. Nat. Commun. 12, 1502 (2021).

39. Iltis, C. et al. A ganglioside-based immune checkpoint enables senescent cells to evade immunosurveillance during aging. Nat. Aging 1–18 (2024) doi:10.1038/s43587-024-00776-z.

40. Ryu, S. et al. Siglec-F–expressing neutrophils are essential for creating a profibrotic microenvironment in renal fibrosis. J. Clin. Invest. 132, e156876.

41. Engblom, C. et al. Osteoblasts remotely supply lung tumors with cancer-promoting SiglecFhigh neutrophils. Science 358, eaal5081 (2017).

42. Pfirschke, C. et al. Tumor-Promoting Ly-6G+ SiglecFhigh Cells Are Mature and Long-Lived Neutrophils. Cell Rep. 32, 108164 (2020).

43. Feng, Y. et al. Breast cancer development and progression: Risk factors, cancer stem cells, signaling pathways, genomics, and molecular pathogenesis. Genes Dis. 5, 77–106 (2018).

44. Collin, G., Huna, A., Warnier, M., Flaman, J.-M. & Bernard, D. Transcriptional repression of DNA repair genes is a hallmark and a cause of cellular senescence. Cell Death Dis. 9, 259 (2018).

45. Saul, D. et al. A new gene set identifies senescent cells and predicts senescence-associated pathways across tissues. Nat. Commun. 13, 4827 (2022).

46. Fridman, A. & Tainsky, M. Critical pathways in cellular senescence and immortalization revealed by gene expression profiling. Oncogene 27, 10.1038/onc.2008.213 (2008).

47. Jones, A. B., Rocco, A., Lamb, L. S., Friedman, G. K. & Hjelmeland, A. B. Regulation of NKG2D Stress Ligands and Its Relevance in Cancer Progression. Cancers 14, 2339 (2022).

48. Vanacker, H. et al. Emerging Role of the Unfolded Protein Response in Tumor Immunosurveillance. Trends Cancer 3, 491–505 (2017).

49. Masuda, H. et al. Role of Epidermal Growth Factor Receptor in Breast Cancer. Breast Cancer Res. Treat. 136, 10.1007/s10549-012-2289–9 (2012).

50. Sparta, B. et al. Receptor Level Mechanisms Are Required for Epidermal Growth Factor (EGF)-stimulated Extracellular Signal-regulated Kinase (ERK) Activity Pulses. J. Biol. Chem. 290, 24784–24792 (2015).

51. Hamanaka, R. B., Bennett, B. S., Cullinan, S. B. & Diehl, J. A. PERK and GCN2 Contribute to eIF2α Phosphorylation and Cell Cycle Arrest after Activation of the Unfolded Protein Response Pathway. Mol. Biol. Cell 16, 5493–5501 (2005).

52. Huber, A.-L. et al. p58IPK-Mediated Attenuation of the Proapoptotic PERK-CHOP Pathway Allows Malignant Progression upon Low Glucose. Mol. Cell 49, 1049–1059 (2013).

53. Holicek, P., et al. Type I interferon and cancer. Immunol. Rev. 321, 115–127 (2024).

54. Iannello, A., Thompson, T. W., Ardolino, M., Lowe, S. W. & Raulet, D. H. p53-dependent chemokine production by senescent tumor cells supports NKG2D-dependent tumor elimination by natural killer cells. J. Exp. Med. 210, 2057–2069 (2013).

55. Coelho, M. A. et al. Oncogenic RAS Signaling Promotes Tumor Immunoresistance by Stabilizing PD-L1 mRNA. Immunity 47, 1083–1099.e6 (2017).

56. Wang, T.-W. et al. Blocking PD-L1–PD-1 improves senescence surveillance and ageing phenotypes. Nature 611, 358–364 (2022).

57. Logue, S. E. et al. Inhibition of IRE1 RNase activity modulates the tumor cell secretome and enhances response to chemotherapy. Nat. Commun. 9, 3267 (2018).

58. Lhomond, S., et al. Dual IRE1 RNase functions dictate glioblastoma development. EMBO Mol. Med. 10, e7929 (2018).

59. Hosomi, S. et al. Intestinal epithelial cell endoplasmic reticulum stress promotes MULT1 up-regulation and NKG2D-mediated inflammation. J. Exp. Med. 214, 2985–2997 (2017).

60. Martins, I. et al. Restoration of the immunogenicity of cisplatin-induced cancer cell death by endoplasmic reticulum stress. Oncogene 30, 1147–1158 (2011).

61. Senovilla, L. et al. An Immunosurveillance Mechanism Controls Cancer Cell Ploidy. Science 337, 1678–1684 (2012).

62. Coppé, J.-P. et al. Senescence-Associated Secretory Phenotypes Reveal Cell-Nonautonomous Functions of Oncogenic RAS and the p53 Tumor Suppressor. PLOS Biol. 6, e301 (2008).

63. Catanzaro, J. M. et al. Oncogenic Ras induces inflammatory cytokine production by up-regulating the squamous cell carcinoma antigens SerpinB3/B4. Nat. Commun. 5, 3729 (2014).

64. Hamarsheh, S., Groß, O., Brummer, T. & Zeiser, R. Immune modulatory effects of oncogenic KRAS in cancer. Nat. Commun. 11, 5439 (2020).

65. Xue, W. et al. Senescence and tumour clearance is triggered by p53 restoration in murine liver carcinomas. Nature 445, 656–660 (2007).

66. Kang, T.-W. et al. Senescence surveillance of pre-malignant hepatocytes limits liver cancer development. Nature 479, 547–551 (2011).

67. Abbas, T. & Dutta, A. p21 in cancer: intricate networks and multiple activities. Nat. Rev. Cancer 9, 400 (2009).

68. Sturmlechner, I. et al. p21 produces a bioactive secretome that places stressed cells under immunosurveillance. Science 374, eabb3420 (2021).

69. Onorati, A. et al. Upregulation of PD-L1 in Senescence and Aging. Mol. Cell. Biol. 42, e00171 (2022).

70. Majewska, J. et al. p16-dependent increase of PD-L1 stability regulates immunosurveillance of senescent cells. Nat. Cell Biol. 26, 1336–1345 (2024).

71. Yi, M., Niu, M., Xu, L., Luo, S. & Wu, K. Regulation of PD-L1 expression in the tumor microenvironment. J. Hematol. Oncol.J Hematol Oncol 14, 10 (2021).

72. Jiang, X. et al. Role of the tumor microenvironment in PD-L1/PD-1-mediated tumor immune escape. Mol. Cancer 18, 10 (2019).

73. Niu, C. et al. PD-1-positive Natural Killer Cells have a weaker antitumor function than that of PD-1-negative Natural Killer Cells in Lung Cancer. Int. J. Med. Sci. 17, 1964 (2020).

74. Batista, A. et al. IRE1α regulates macrophage polarization, PD-L1 expression, and tumor survival. PLoS Biol. 18, e3000687 (2020).

75. Rolas, L. et al. Senescent endothelial cells promote pathogenic neutrophil trafficking in inflamed tissues. EMBO Rep. 1–28 (2024) doi:10.1038/s44319-024-00182-x.

76. Cambier, S., Gouwy, M. & Proost, P. The chemokines CXCL8 and CXCL12: molecular and functional properties, role in disease and efforts towards pharmacological intervention. Cell. Mol. Immunol. 20, 217–251 (2023).

77. Raulet, D. H. & Guerra, N. Oncogenic stress sensed by the immune system: role of NK cell receptors. Nat. Rev. Immunol. 9, 568–580 (2009).

78. Matlung, H. L. et al. Neutrophils Kill Antibody-Opsonized Cancer Cells by Trogoptosis. Cell Rep. 23, 3946–3959.e6 (2018).

79. Tecchio, C. & Cassatella, M. A. Neutrophil-derived chemokines on the road to immunity. Semin. Immunol. 28, 119–128 (2016).

80. Sagiv, A. et al. NKG2D ligands mediate immunosurveillance of senescent cells. Aging 8, 328–344 (2016).

81. Guia, S. et al. Genome-wide CRISPR/Cas9 screen reveals factors that influence the susceptibility of tumor cells to NK cell-mediated killing. 2024.10.08.615667 Preprint at 10.1101/2024.10.08.615667 (2024).

82. Guglietta, S. et al. Coagulation induced by C3aR-dependent NETosis drives protumorigenic neutrophils during small intestinal tumorigenesis. Nat. Commun. 7, 11037 (2016).

