## Supplementary Table 1 for "Early Neutrophil Sensing Shapes the Innate Response to Preneoplastic Mammary Cells"

**Table S1 : Differentially Expressed Genes eHMEC vs Ctl HMEC**

| SYMBOL | Localisation | logFC | adj.P.Val |
|---|---|---|---|
| ANOS1 | Both | 1,028078501 | 9,78645E-06 |
| HBEGF | Both | 2,039493576 | 5,23097E-05 |
| TGFA | Both | 0,829517085 | 0,000187767 |
| ANTXR2 | Both | 0,780731708 | 0,000379555 |
| PLAUR | Both | 1,355860675 | 0,001666771 |
| CD55 | Both | 0,683767936 | 0,005196163 |
| IL1RL1 | Both | 1,429518289 | 0,006419643 |
| DSCAM | Both | 0,697797156 | 0,008041548 |
| IL1R2 | Both | 1,214516047 | 0,015381753 |
| HPGD | Other | 2,784895824 | 1,23507E-07 |
| BIRC3 | Other | 2,022755045 | 5,45531E-06 |
| LCP1 | Other | 1,020686147 | 9,78645E-06 |
| CHAC1 | Other | 1,034064127 | 1,0157E-05 |
| TAGLN3 | Other | 1,821813311 | 1,20614E-05 |
| ARHGAP22 | Other | 0,895464307 | 1,36371E-05 |
| ETS1 | Other | 0,74796164 | 2,19633E-05 |
| NEDD4L | Other | 0,780071666 | 2,84324E-05 |
| PMAIP1 | Other | 0,910462146 | 3,51168E-05 |
| HERPUD1 | Other | 0,860441407 | 4,10625E-05 |
| UCA1 | Other | 0,814724838 | 4,54001E-05 |
| MAFF | Other | 0,946715782 | 4,9331E-05 |
| KRT8 | Other | 0,821924665 | 5,36037E-05 |
| CTH | Other | 1,238876245 | 0,000102285 |
| DNAJB9 | Other | 0,650247556 | 0,00011187 |
| PDE1C | Other | 1,169246897 | 0,000120164 |
| ASNS | Other | 0,598052288 | 0,000139474 |
| CBS | Other | 0,695213851 | 0,000141363 |
| PHLDA1 | Other | 0,636412706 | 0,0001525 |
| NFKBIE | Other | 0,668687103 | 0,000209999 |
| KCTD12 | Other | 0,62729369 | 0,000223677 |
| NAV3 | Other | 0,822012233 | 0,000246574 |
| KRTAP2-3 | Other | 1,643842298 | 0,000249111 |
| DDX58 | Other | 0,754514149 | 0,000277069 |
| UGCG | Other | 0,762568562 | 0,000289277 |
| PPP1R3B | Other | 0,682775185 | 0,000289277 |
| CSGALNACT2 | Other | 0,624933397 | 0,000289277 |
| BIK | Other | 0,61580067 | 0,000289277 |
| TRIB3 | Other | 0,613220437 | 0,000289277 |
| KLF6 | Other | 0,580423233 | 0,000289277 |
| PLOD2 | Other | 0,632021665 | 0,000309211 |
| TNIP1 | Other | 0,900058121 | 0,000318888 |
| CPEB4 | Other | 0,861448883 | 0,000379555 |
| ARNTL2 | Other | 0,79033392 | 0,000403701 |
| TRIB1 | Other | 0,761876925 | 0,000415696 |
| PPIF | Other | 0,676090712 | 0,000418065 |
| CCNA1 | Other | 1,111204822 | 0,000471228 |
| FAM214B | Other | 0,851210064 | 0,000474984 |

| | | | |
|---|---|---|---|
| FAM110C | Other | 0,857464974 | 0,000515454 |
| TNFAIP3 | Other | 1,567084675 | 0,000545185 |
| TXNRD1 | Other | 0,633556711 | 0,000594655 |
| GFPT1 | Other | 0,642850012 | 0,000603925 |
| EGR1 | Other | 1,06136785 | 0,00068359 |
| KDM7A | Other | 0,691168619 | 0,00072695 |
| MAOA | Other | 0,929461794 | 0,000759963 |
| TRIM8 | Other | 0,645160516 | 0,000759963 |
| PTGS2 | Other | 2,094166408 | 0,000855466 |
| PCK2 | Other | 0,66643698 | 0,000901983 |
| KRT34 | Other | 1,603903237 | 0,000978344 |
| PLCB4 | Other | 0,627369612 | 0,00098832 |
| INSIG2 | Other | 0,646901179 | 0,001025441 |
| GSAP | Other | 0,588156915 | 0,00112556 |
| ROBO4 | Other | 0,669694991 | 0,00140697 |
| ISG20 | Other | 1,544426108 | 0,001409188 |
| FAM83A | Other | 0,630207936 | 0,001417369 |
| MARCHF4 | Other | 0,591523272 | 0,001491869 |
| CORO1A | Other | 0,622694303 | 0,001512626 |
| GRAMD2B | Other | 0,695862647 | 0,001534493 |
| NABP1 | Other | 0,649289522 | 0,001537799 |
| CYB5R2 | Other | 0,902229304 | 0,001542961 |
| SOCS1 | Other | 0,620024183 | 0,001597626 |
| MAP1B | Other | 0,696342334 | 0,001885569 |
| USP43 | Other | 0,778002212 | 0,00242875 |
| HMOX1 | Other | 1,052067819 | 0,002440657 |
| DUSP5 | Other | 0,796046227 | 0,002523522 |
| EHF | Other | 1,197266572 | 0,003061926 |
| LCE1F | Other | 0,893635046 | 0,003061926 |
| IER2 | Other | 0,664515851 | 0,003182398 |
| DNTTIP1 | Other | 0,63488226 | 0,00326767 |
| CHST2 | Other | 0,911010085 | 0,003388071 |
| RASA2 | Other | 0,592192295 | 0,003464578 |
| BHLHE40 | Other | 0,585997091 | 0,003811432 |
| HS3ST2 | Other | 0,590002034 | 0,004275777 |
| ZNF114 | Other | 0,954718715 | 0,004763535 |
| MAP1S | Other | 0,580475495 | 0,004988813 |
| KCTD11 | Other | 0,665188046 | 0,004995433 |
| ARID3B | Other | 0,633917059 | 0,005514545 |
| NFKBIZ | Other | 0,992131886 | 0,005811143 |
| TICAM1 | Other | 0,688802179 | 0,006291324 |
| PPP1R15A | Other | 0,896880066 | 0,006419643 |
| ZC3H12C | Other | 0,599645863 | 0,006969537 |
| GLRX | Other | 1,398911484 | 0,008517688 |
| ITPKC | Other | 0,778328883 | 0,008925873 |
| DUSP1 | Other | 0,708714553 | 0,009236685 |
| CREBRF | Other | 0,643504353 | 0,010214291 |
| NFKB2 | Other | 0,623383581 | 0,011714589 |
| ABL2 | Other | 0,655651446 | 0,013385725 |
| PRDM1 | Other | 1,132420325 | 0,017841929 |

| | | | |
|---|---|---|---|
| CGN | Other | 0,718668037 | 0,025616806 |
| HSD17B2 | Other | 1,093435253 | 0,027973478 |
| KRT33B | Other | 0,702228961 | 0,034045768 |
| SPRY2 | Other | 0,862304808 | 0,037290465 |
| CRCT1 | Other | 1,964413268 | 0,039387831 |
| TMEM200A | Plasma membrane | 1,298082266 | 5,45531E-06 |
| S1PR1 | Plasma membrane | 1,160759492 | 9,78645E-06 |
| AKAP12 | Plasma membrane | 1,130170534 | 9,78645E-06 |
| COL13A1 | Plasma membrane | 0,8750619 | 3,86297E-05 |
| VLDLR | Plasma membrane | 0,772915264 | 4,9331E-05 |
| ERRFI1 | Plasma membrane | 0,707392142 | 6,49893E-05 |
| MGLL | Plasma membrane | 1,117214562 | 6,53575E-05 |
| ADAM8 | Plasma membrane | 0,672419582 | 6,58399E-05 |
| FAM241B | Plasma membrane | 0,650886268 | 9,89821E-05 |
| SLC41A2 | Plasma membrane | 0,732168353 | 0,000148554 |
| MYADM | Plasma membrane | 0,594063555 | 0,0001525 |
| EPHA2 | Plasma membrane | 0,587748297 | 0,0001525 |
| SMURF2 | Plasma membrane | 0,777759586 | 0,000185505 |
| SEMA7A | Plasma membrane | 0,658703528 | 0,000187767 |
| PHLDA2 | Plasma membrane | 0,697259114 | 0,000246574 |
| CD274 | Plasma membrane | 0,753068313 | 0,000262909 |
| DAB2 | Plasma membrane | 0,738017101 | 0,000289277 |
| STYK1 | Plasma membrane | 0,65655089 | 0,000289277 |
| CHIC2 | Plasma membrane | 0,787399742 | 0,000379555 |
| PTPRE | Plasma membrane | 0,591964918 | 0,000491691 |
| IL13RA2 | Plasma membrane | 2,47417114 | 0,000521127 |
| TMEFF1 | Plasma membrane | 0,867447257 | 0,000673744 |
| RAB3B | Plasma membrane | 0,706802594 | 0,000775089 |
| ICAM1 | Plasma membrane | 1,32576503 | 0,000823109 |
| ITGA2 | Plasma membrane | 0,833633424 | 0,000859444 |
| TM4SF19 | Plasma membrane | 1,412111959 | 0,002097483 |
| CYSTM1 | Plasma membrane | 0,600655562 | 0,002104432 |
| DLG1 | Plasma membrane | 0,659533532 | 0,002377826 |
| HAS2 | Plasma membrane | 1,769117088 | 0,002818594 |
| PARD6B | Plasma membrane | 0,908631179 | 0,003315766 |
| TJP2 | Plasma membrane | 0,629521507 | 0,003464578 |
| IER3 | Plasma membrane | 0,738886959 | 0,003617695 |
| TRPV3 | Plasma membrane | 0,627433785 | 0,003835108 |
| BNIP3L | Plasma membrane | 0,810356669 | 0,004680796 |
| RALA | Plasma membrane | 0,728618392 | 0,004763535 |
| OCLN | Plasma membrane | 1,035835638 | 0,005179173 |
| CD200 | Plasma membrane | 1,051218079 | 0,005582132 |
| RALGPS2 | Plasma membrane | 0,736999626 | 0,006161819 |
| TNFSF15 | Plasma membrane | 0,867032955 | 0,006419643 |
| GPRC5A | Plasma membrane | 1,070072328 | 0,006834912 |
| OXTR | Plasma membrane | 0,776888537 | 0,008117986 |
| PLD5 | Plasma membrane | 0,753929524 | 0,008586049 |
| ZP4 | Plasma membrane | 0,601702692 | 0,008707311 |
| SLC2A3 | Plasma membrane | 0,748945581 | 0,013173296 |
| SLC16A4 | Plasma membrane | 0,740994045 | 0,015786679 |

| | | | |
|---|---|---|---|
| GJB4 | Plasma membrane | 0,611141869 | 0,027083673 |
| STC2 | Secreted | 1,001797952 | 5,45531E-06 |
| CXCL3 | Secreted | 2,444318525 | 1,30977E-05 |
| CXCL8 | Secreted | 2,966995294 | 1,36371E-05 |
| IL1A | Secreted | 1,21843582 | 1,6549E-05 |
| CXCL1 | Secreted | 1,849479886 | 3,21236E-05 |
| PLAT | Secreted | 0,727128223 | 5,36037E-05 |
| INHBA | Secreted | 1,603058906 | 0,000141076 |
| ADAMTS1 | Secreted | 1,525709167 | 0,0001525 |
| EDN1 | Secreted | 1,266497495 | 0,000289277 |
| NLRP3 | Secreted | 1,104301824 | 0,000289277 |
| PLAU | Secreted | 0,943649514 | 0,000289277 |
| ADAMTS6 | Secreted | 0,589497037 | 0,000289277 |
| LAMB3 | Secreted | 0,587013845 | 0,000289277 |
| CLCF1 | Secreted | 0,956902716 | 0,000472543 |
| PDZD2 | Secreted | 0,62856683 | 0,000488909 |
| CXCL2 | Secreted | 1,635886729 | 0,000624424 |
| RNASE7 | Secreted | 0,825984384 | 0,000902137 |
| CST6 | Secreted | 0,977726428 | 0,001095039 |
| SRGN | Secreted | 0,711935753 | 0,00112556 |
| CSF2 | Secreted | 2,121380521 | 0,001674137 |
| LAMC2 | Secreted | 0,692656637 | 0,001674137 |
| LAMA3 | Secreted | 0,599326693 | 0,002729173 |
| LIF | Secreted | 0,920530202 | 0,004275777 |
| IL24 | Secreted | 0,591814504 | 0,005799342 |
| PPBP | Secreted | 0,922802548 | 0,008041548 |
| DEFB103B | Secreted | 1,352415983 | 0,009762495 |
| ESM1 | Secreted | 1,416896142 | 0,011253721 |
| ACP7 | Secreted | 0,63930806 | 0,012171695 |
| SCG5 | Secreted | 0,916434899 | 0,012680633 |
| IL32 | Secreted | 0,587872481 | 0,017881134 |
| SERPINB2 | Secreted | 1,126493295 | 0,025099838 |
| IL1B | Secreted | 0,766648963 | 0,031346396 |
| BMP2 | Secreted | 0,833541418 | 0,037174027 |
| SERPINB1 | Secreted | 0,777576517 | 0,039956203 |
| STC1 | Secreted | 0,684128159 | 0,041744407 |
