## Supplementary Tables 2-7 for "Early Neutrophil Sensing Shapes the Innate Response to Preneoplastic Mammary Cells"

### Table S2: Panels of antibodies for cytometry assays

| a. Antibodies for Flow cytometry BRCA1 model | Manufacturer | Reference | Clone | Dilution |
|---|---|---|---|---|
| BV711 anti-mouse CD8a | Biolegend | 100747 | 53-6.7 | 1:200 |
| BV650 anti-mouse CD3 | BD | 564378 | 145-2C11 | 1:50 |
| Zombie Aqua | Biolegend | 423102 | | 1:400 |
| V450 anti-mouse CD19 | BD | 560376 | 1D3 | 1:50 |
| FITC anti-mouse Ly6G | BD | 551460 | 1A8 | 1:100 |
| PE anti-mouse Siglec F | BD | 552126 | E50-2440 | 1:100 |
| PE-Cy7 anti-mouse FoxP3 | eBioscience | 25-5773-82 | FJK-16s | 1:100 |
| PE-Cy5 anti-mouse CD4 | BD | 553654 | H129.19 | 1:200 |
| PE-Dazzle 594 anti-mouse CD64 | Biolegend | 139319 | X54-5/7.1 | 1:200 |
| APC anti-mouse CD49b | BD | 560628 | DX5 | 1:50 |
| Alexa Fluor 700 anti-mouse CD45 | Biolegend | 103128 | 30-F11 | 1:400 |

| b. Antibodies for cell sorting of Epithelial Cells for MMTV-Neu model | Source | Reference | Clone | Dilution |
|---|---|---|---|---|
| Viability, DAPI | | | | 1X |
| BV605 anti-mouse TER-119 | BD | 747740 | Clone TER-119 | 1:200 |
| FITC anti-mouse Ly6G | BD | 551460 | 1A8 | 1:400 |
| PE anti-mouse Siglec F | BD | 552126 | E50-2440 | 1:100 |
| PE-Cy7 anti-mouse EpCAM | Biolegend | 118216 | G8.8 | 1:200 |
| Alexa Fluor 700 anti-mouse CD45 | Biolegend | 103128 | 30-F11 | 1:400 |
| APC-Cy7 anti-mouse Ly6C | Biolegend | 560596 | AL-21 | 1:200 |

| c. Antibodies for Cell sorting of Neutrophils for MMTV-Neu model | Manufacturer | Reference | Clone | Dilution |
|---|---|---|---|---|
| Purified anti-mouse CD16/32 | Biolegend | 101302 | 93 | 1:50 |
| FITC anti-mouse Ly6G | BD | 551460 | 1A8 | 1:400 |
| PE anti-mouse Siglec F | BD | 552126 | E50-2440 | 1:100 |
| PE-Cy7 anti-mouse CD326 | Biolegend | 118216 | G8.8 | 1:200 |
| APC anti-mouse CD3 | Biolegend | 100236 | 17A2 | 1:100 |
| APC anti-mouse CD4 | Biolegend | 100412 | GK1.5 | 1:400 |
| APC anti-mouse CD8a | Biolegend | 100712 | 53-6-7 | 1:200 |
| APC anti-mouse CD11c | BD | 550261 | HL3 | 1:100 |
| APC anti-mouse CD19 | Biolegend | 115512 | 6D5 | 1:200 |
| APC anti-mouse CD335 | Biolegend | 137608 | 29A1.4 | 1:100 |
| APC anti-mouse TER-119 | Biolegend | 116212 | TER-119 | 1:200 |
| Alexa Fluor 700 anti-mouse CD45.2 | Biolegend | 109822 | 104 | 1:400 |
| APC-Cy7 anti-mouse Ly6C | BD | 560596 | Al-21 | 1:200 |

**d. Antibodies for Immune landscape of MMTV-Neu model**

| | | | | | |
|---|---|---|---|---|---|
| CD3 | CD19 | CD45 | CD172a/SIRPa | IA-IE | Siglec F |
| CD4 | CD25 | CD62L | B220 | IgD | Siglec H |
| CD8 | CD27 | CD64 | DX5/CD49d | IgM | TCRgd |
| CD11b | CD38 | CD103 | EpCAM | Ly6C | XCR1 |
| CD11c | CD44 | CD127 | FoxP3 | Ly6G | LiveDead Blue |

| e. Antibodies for Transwell Migration Assay (Isolated Neutrophils) | Manufacturer | Reference | Clone | Dilution |
|---|---|---|---|---|
| BV421 anti-human CD66b | BD | 562940 | G10F5 | 1:30 |
| PE-CF594 anti-human CD15 | BD | 562485 | MC480 | 1:50 |
| Alexa Fluor 700 anti-human CD45 | BD | 560566 | HI30 | 1:25 |
| Zombie NIR | Biolegend | 423106 | | 1:800 |

| f. Antibodies for Transwell Migration Assay (Total immune cells and PBMCs) | Manufacturer | Reference | Clone | Dilution |
|---|---|---|---|---|
| BUV395 anti-human CD45 | BD | 563792 | HI30 | 1:50 |
| Live/Dead Blue | Thermofisher | L34962 | | 1:500 |
| BUV737 anti-human CD89 | BD | 749244 | A59 | 1:20 |
| BUV805 anti-human CD15 | BD | 742057 | W6D3 | 1:20 |
| BV421 anti-human CD3 | Biolegend | 300434 | UCHT1 | 1:50 |
| BV711 anti-human CD4 | BD | 563028 | SK3 | 1:50 |
| BV750 anti-human CD56 | Biolegend | 362556 | 5.1H11 | 1:20 |
| BB515 anti-human CD20 | BD | 564568 | 2H7 | 1:100 |
| PerCP-Cy5.5 anti-human CD11c | Biolegend | 337210 | Bu15 | 1:50 |
| PE anti-human CD66b | Miltenyi | 130-104-396 | REA 306 | 1:100 |
| PE-Dazzle 594 anti-human CD123 | Biolegend | 306034 | 6H6 | 1:40 |
| PE-Cy7 anti-human CD19 | BD | 557835 | SJ25C1 | 1:50 |
| APC anti-human CD14 | BD | 555399 | M5E2 | 1:25 |
| Alexa Fluor 700 anti-human CD8 | BD | 557945 | RPA-T8 | 1:25 |
| APC-H7 anti-human HLA-DR | BD | 561358 | G46-6 | 1:50 |

| g. Antibodies for p-STAT5 Activation | Manufacturer | Reference | Clone | Dilution |
|---|---|---|---|---|
| BV421 anti-human CD66b | BD | 562940 | G10F5 | 1:30 |
| V500 anti-human CD15 | BD | 561585 | HI98 | 1:50 |
| BV711 anti-human CD14 | Biolegend | 301838 | M5E2 | 1:25 |
| PE-CF594 anti-human p-STAT5 | BD | 562501 | 47 | 1:100 |
| APC anti-human Siglec8 | Miltenyi | 130-117-975 | REA1045 | 1:50 |
| Alexa Fluor 700 anti-human CD45 | BD | 560566 | HI30 | 1:25 |
| Zombie NIR | Biolegend | 423106 | | 1:800 |

| h. Antibodies for ROS Production | Manufacturer | Reference | Clone | Dilution |
|---|---|---|---|---|
| BV421 anti-human CD66b | BD | 562940 | G10F5 | 1:30 |
| PE-CF594 anti-human CD15 | BD | 562485 | MC480 | 1:50 |
| Alexa Fluor 700 anti-human CD45 | BD | 560566 | HI30 | 1:25 |
| Zombie NIR | Biolegend | 423106 | | 1:800 |

| i. Antibodies for Co-culture Assay | Manufacturer | Reference | Clone | Dilution |
|---|---|---|---|---|
| BV421 anti-human CD18 | BD | 743370 | 6.7 | 1:20 |
| BV711 anti-human CD11b | BD | 740771 | ICRF4 | 1:20 |
| FITC anti-human CD66b | BD | 555724 | G10F5 | 1:40 |
| PE anti-human ULBP2-5-6 | R&D | FAB1298P | 165903 | 1:2,5 |
| PE-CF594 anti-human CD63 | BD | 565403 | H5C6 | 1:50 |
| PE-Cy7 anti-human PD-L1 | Biolegend | 329718 | 29E2A3 | 1:50 |
| APC anti-human ICAM1 | Miltenyi | 130-121-342 | REA266 | 1:11 |
| Alexa Fluor 700 anti-human CD45 | BD | 560566 | HI30 | 1:25 |
| Zombie NIR | Biolegend | 423106 | | 1:800 |

| j. Antibodies for *In Vivo* Matrigel Plug Assay | Manufacturer | Reference | Clone | Dilution |
|---|---|---|---|---|
| BUV395 anti-mouse CD11b | BD | 563553 | M1/70 | 1:100 |
| BV421 anti-mouse PD-L1 | Biolegend | 124315 | 10F.9G2 | 1:100 |
| BV605 anti-mouse CD54 | BD | 740343 | 3E2 | 1:200 |
| BV786 anti-mouse CD62L | BD | 564109 | MEL-14 | 1:400 |
| Vio B515 anti-mouse CD107a | Miltenyi | 130-111-320 | REA777 | 1:100 |
| PerCP vio 770 anti-mouse Ly6G | Miltenyi | 130-117-500 | REA526 | 1:50 |
| PE anti-mouse SiglecF | BD | 552126 | E50-2440 | 1:100 |
| PE-Dazzle 594 anti-mouse CD3 | Biolegend | 100348 | 145-2C11 | 1:25 |
| PE-Dazzle 594 anti-mouse CD19 | Biolegend | 115554 | 6D5 | 1:10 |
| PE-Dazzle 594 anti-mouse CD64 | Biolegend | 139319 | X54-5/7.1 | 1:100 |
| PE-Vio 770 anti-mouse CD69 | Miltenyi | 130-115-577 | REA937 | 1:100 |
| APC anti-mouse NK1.1 | Biolegend | 108710 | PK136 | 1:100 |
| Alexa Fluor 700 anti-mouse CD45 | Biolegend | 103128 | 30-F11 | 1:200 |
| ViaKrome 808 | Beckman | C36628 | | 1:40 |

**Table S3 : Gene sets and signatures**

| **a.** ssGSEA analysis Geneset (Mouse models) | Species | Collection |
|---|---|---|
| HALLMARK_UNFOLDED_PROTEIN_RESPONSE | Mouse | MH |
| HALLMARK_REACTIVE_OXYGEN_SPECIES_PATHWAY | Mouse | MH |
| HALLMARK_HYPOXIA | Mouse | MH |
| HALLMARK_GLYCOLYSIS | Mouse | MH |
| HALLMARK_TGF_BETA_SIGNALING | Mouse | MH |
| REACTOME_UNFOLDED_PROTEIN_RESPONSE_UPR | Mouse | M2 |
| GOBP_ENDOPLASMIC_RETICULUM_UNFOLDED_PROTEIN_RESPONSE | Mouse | M5 |
| GOBP_ERBB2_SIGNALING_PATHWAY | Mouse | M5 |
| GOBP_RESPONSE_TO_ENDOPLASMIC_RETICULUM_STRESS | Mouse | M5 |
| GOBP_NEUTROPHIL_ACTIVATION | Mouse | M5 |
| GOBP_NEUTROPHIL_MEDIATED_IMMUNITY | Mouse | M5 |
| GOBP_NEUTROPHIL_DEGRANULATION | Mouse | M5 |
| GOBP_CYTOPLASMIC_TRANSLATION | Mouse | M5 |
| GOBP_PROTEIN8FOLDING | Mouse | M5 |
| GOBP_REGULATION_OF_CELL_PROLIFERATION | Mouse | M5 |
| GOBP_EPITHELIAL_CELL_PROLIFERATION | Mouse | M5 |
| GOBP_POSITIVE_REGULATION_OF_EPITHELIAL_CELL_PROLIFERATION | Mouse | M5 |
| GOBP_REGULATION_OF_IMMUNE_RESPONSE | Mouse | M5 |
| GOBP_ACTIVATION_OF_IMMUNE_RESPONSE | Mouse | M5 |
| GOBP_GRANULOCYTE_MIGRATION | Mouse | M5 |
| GOBP_GRANULOCYTE_CHEMOTAXIS | Mouse | M5 |
| GOBP_ACTIVATION_OF_INNATE_IMMUNE_RESPONSE | Mouse | M5 |

| **b.** ssGSEA analysis publicly dataset (Mouse models) | Species | Genes |
|---|---|---|
| mN1_signature | Mouse | Chsy1 ; Retnlb ; Mmp8 ; Scnn1a ; Dhrs7 ; Retnlg |
| mN5_signature | Mouse | Gpnmb ; Gstm1 ; Ccl3 ; Psap ; Rab15 ; Fnip2 ; Cstb ; Mreg ; Ctsb ; Rnh1 ; Hilpda ; Itpr2 ; Gns ; Cd63 ; Atp6v1c1 |
| mNeu08_signature | Mouse | Mmp8 ; Mmp9 ; Camp ; Ngp ; Lcn2 |
| mNeu12_signature | Mouse | Ccl3 ; Ccl4 |
| T2_signature | Mouse | Cd300ld ; Cxcr2 ; Dusp1 ; Gbp2 ; Ifitm1 ; Il1b ; Isg15 ; Jaml ; Junb ; Msrb1 ; Osm ; S100a6 ; Selplg ; Slpi |
| T3_signature | Mouse | Atf3 ; Ccl3 ; Ccl4 ; Cd274 ; Cstb ; Cxcl3 ; Hcar2 ; Hilpda ; Hk2 ; Hmox1 ; Ier3 ; Jun ; Plin2 ; Spp1 ; Tgif1 ; Tnfrsf23 ; Vegfa ; Zeb2 ; Ldha ; Mif |
| SiglecF_high_vs_SiglecF_low | Mouse | Upregulated : Adam10 ; Adam17 ; Ccl3 ; Ccl4 ; Cd24a ; Cd63 ; Cd274 ; Csf1 ; Csf2rb2 ; Ctsb ; E2f4 ; Fcgr2b ; Ffar2 ; Havcr2 ; Hif1a ; Il1a ; Itgax ; Siglecf ; Scimp ; Spp1 ; Tnf ; Vegfa ; Xbp1<br>Downregulated : Anxa1 ; Anxa2 ; Cd244a ; Fas ; Foxp1 ; Il15 ; Irf1 ; Ly6g ; H2-D1 ; H2-Q7 ; H2-Q10 ; Itgal ; Lrg1 ; Mmp8 ; Timp2 |

| **c.** ssGSEA analysis Geneset (HMEC model) | Species | Collection |
|---|---|---|
| HALLMARK_DNA_REPAIR | Human | H |
| BILD_HRAS_ONCOGENIC_SIGNATURE | Human | C2-CPG |
| FRIDMAN_SENESCENCE_UP | Human | C2-CPG |
| SAUL_SEN_MAYO | Human | C2-CPG |
| GOBP_ATF6_MEDIATED_UNFOLDED_PROTEIN_RESPONSE | Human | C5 |
| GOBP_IRE1_MEDIATED_UNFOLDED_PROTEIN_RESPONSE | Human | C5 |
| GOBP_PERK_MEDIATED_UNFOLDED_PROTEIN_RESPONSE | Human | C5 |
| GOBP_REGULATION_OF_RESPONSE_TO_STRESS | Human | C5 |
| GOBP_RESPONSE_TO_ENDOPLASMIC_RETICULUM_STRESS | Human | C5 |
| GOBP_RESPONSE_TO_TYPE_I_INTERFERON | Human | C5 |

### Table S4 : Panel of antibodies for Multiplex Immunofluorescence

| Antibodies for mIF | Manufacturer | Reference | Clone | Dilution |
|---|---|---|---|---|
| rabbit polyclonal antibody anti-EpCAM | Abcam | ab71916 | polyclonal | 1:500 |
| rabbit anti-Mouse Ly6G | CST | 87048S | E6Z1T | 1:300 |
| rabbit anti-Mouse CD45 | CST | 70257 | D3F8Q | 1:300 |

### Table S5 : Kits ECLIA and ELISA

| a. Kits ECLIA | Manufacturer | Reference |
|---|---|---|
| U-Plex Custom Biomarker Group I (Ms) Assay | Meso Scale Discovery® | K15069M-1 |
| U-Plex TGF-b Combo (mouse) | Meso Scale Discovery® | K15242K-1 |

| b. Kits ELISA | Manufacturer | Reference |
|---|---|---|
| Human GM-CSF DuoSet ELISA | R&D Systems™ - Bio-Techne | DY215 |
| Human CXCL1/GRO alpha DuoSet ELISA | R&D Systems™ - Bio-Techne | DY275 |
| Human G-CSF DuoSet ELISA | R&D Systems™ - Bio-Techne | DY214 |
| Human CXCL2/GRO beta DuoSet ELISA | R&D Systems™ - Bio-Techne | DY276 |
| Human GROγ/CXCL3 ELISA Kit (home-made kit)<br>Recombinant human GROγ Cat300-40<br>Biotinylated rabbit anti-human GROγ Cat500-P105Bt | Peprotech | *homemade* |
| Human IL-8 (CXCL8) Standard ABTS ELISA Development Kit | Peprotech | 900-K18 |

### Table S6 : Antibodies for western blotting

| Antibodies for Western Blot | Manufacturer | Reference | Host | Clone | Dilution |
|---|---|---|---|---|---|
| Ras | Santa-Cruz | sc-520 | Rabbit | C-20 | 1:1000 |
| ERK | Cell signaling | 9102 | Rabbit | Polyclonal | 1:1000 |
| p-ERK | Cell signaling | 9101 | Rabbit | Polyclonal | 1:1000 |
| PERK | Cell signaling | 3192S | Rabbit | C33E10 | 1:1000 |
| p-eif2α | Cell signaling | 9722S | Rabbit | Polyclonal | 1:1000 |
| IRE1α | Cell signaling | 3294P | Rabbit | 14C10 | 1:1000 |
| ATF6 | Abcam | ab122897 | Mouse | 1-7 | 1:1000 |
| Cyclin A | Santa-Cruz | sc-751 | Mouse | H-432 | 1:5000 |
| Cycline B1 | Santa-Cruz | sc-245 | Mouse | GNS1 | 1:1000 |
| Cyclin D1 | Santa-Cruz | sc-246 | Mouse | HD11 | 1:1000 |
| Cyclin D3 | Santa-Cruz | sc-182 | Rabbit | C-16 | 1:250 |
| Cyclin E | Santa-Cruz | sc-247 | Mouse | HE12 | 1:500 |
| Cdk1 | Santa-Cruz | sc-54 | Mouse | 17 | 1:500 |
| Cdk2 | Santa-Cruz | sc-163 | Rabbit | M2 | 1:1000 |
| Cdk4 | Santa-Cruz | sc-260 | Rabbit | C-22 | 1:2000 |
| Cdk6 | Santa-Cruz | sc-177 | Rabbit | C-21 | 1:2000 |
| β-actin | MP Biomedicals | 69100 | Mouse | C4 | 1:50000 |
| αTubulin | Sigma-Aldrich | T5168 | Mouse | B-5-1-2 | 1:2500 |

### Table S7 : Primers for RT-qPCR

| Primers for RT-qPCR | Manufacturer | Forward primer | Reverse Primer |
|---|---|---|---|
| GUS primers | Eurogentech | 5’ CGTGGTTGGAGAGCTCATTTGGAA 3’ | 5’ ATTCCCCAGCACTCTCGTCGGT 3’ |
| DNAJB9 primers | Eurogentech | 5’ TCTTAGGTGTGCCAAAATCGG 3’ | 5’ TGTCAGGGTGGTACTTCATGG 3’ |
| HERPUD primers | Eurogentech | 5’ CAAAAATGCCAGAAATCAACG 3’ | 5’ CTGTCCCCGATTAGAACCAG 3’ |
| HRD1 primers | Eurogentech | 5’ CCAGTACCTCACCGTGCTG 3’ | 5’ GCCTCTGAGCTAGGGATGC 3’ |
| XBP1s primers | Eurogentech | 5’ CTGAGTCCGAATCAGGTGCAG 3’ | 5’ ATCCATGGGGAGATGTTCTGG 3’ |
| PPP15R1A primers | Eurogentech | 5’ GCTTCTGGCAGACCGAAC 3’ | 5’ GTAGCCTGATGGGGTGCTT 3’ |
| CHOP primers | Eurogentech | 5’ CAGAGCTGGAACCTGAGGAG 3’ | 5’ TGGATCAGTCTGGAAAAGCA 3’ |
| RIG-I primers | Eurogentech | 5’ AGCTCAGCTTGATGAGGGACA 3’ | 5’ GTCTGGCATCTGGAACACCA 3’ |
| ISG20 primers | Eurogentech | 5’ CACCCCTCAGCACATGGT 3’ | 5’ TGGAAGTCGTGCTTCAGGT 3’ |
| IRF7 primers | Eurogentech | 5’ CAAGTGCAAGGTGTACTGG 3’ | 5’ CAGGTAGATGGTATAGCGTGG 3’ |
| IRF9 primers | Eurogentech | 5’ CATGCAGAACTGCACACTCA 3’ | 5’ CCAATGTCTGAATGGACTGC 3’ |
| SOCS1 primers | Eurogentech | 5’ TTTTCGCCCTTAGCGTGAAGA 3’ | 5' GAGGCAGTCGAAGCTCTCG 3' |
